# Spatial Total RNA Sequencing of Formalin-Fixed Paraffin Embedded Tissues by spRandom-seq

**DOI:** 10.1101/2025.02.19.638981

**Authors:** Yuan Liao, Shunji Zhang, Jiaye Chen, Yu-sheng Chen, Yuexiao Lyu, Bin Zhu, Haide Chen, Yuyi Zhu, Ziye Xu, Yafei Yin, Xushen Xiong, Nan Liu, Hongshan Guo, Yongcheng Wang

## Abstract

Current oligo(dT) primers-based spatial transcriptomic methods are limited to fresh or fresh-frozen samples due to the low efficiency of oligo(dT) primers in capturing RNAs in degraded or microbial samples. Here, we have developed a random primer-based spatial total RNA sequencing (spRandom-seq) technology for simultaneously capturing whole host and microbial RNAs in formalin-fixed paraffin-embedded (FFPE) tissues. spRandom-seq eliminated 3’ or 5’ gene-body biases and outperformed oligo(dT)-based 10X Visium with an 8-fold higher capturing rate for lncRNA and other non-polyadenylated RNA biotypes, including miRNA, snRNA and miscRNA. In the clinical FFPE sections of breast cancer, we revealed the inherent heterogeneity within the tumor region. We also simultaneously captured host and microbial RNAs in *Klebsiella pneumoniae*-infected sections. Totally, spRandom-seq provided a versatile tool for both clinical pathology and infection biology with established spatial platforms, ensuring ease of operation and large-scale applications.

## Introduction

Patients’ formalin-fixed paraffin-embedded (FFPE) tissues are invaluable clinical repositories. However, the total RNAs in FFPE samples are prone to fragmentation and degradation, limiting the utility of oligo-dT primers. The absence of polyadenylated RNA also becomes critical when investigating host-microbial interactions, as pathogenic microorganisms colonize host tissues, and their spatial distribution patterns are linked to disease progression.^1-7^

Current spatially resolved transcriptomics techniques have two major categories: image-based and sequencing-based. Image-based methods such as 10X Xenium, MERFISH^8,9^, seqFISH+^10^ and CosMx^11^ have been applied to FFPE samples, but their targeted probes still limit them and lack full-length gene coverage. The sequencing-based methods rely on poly(A) tails of mRNA, primarily applicable to fresh or fresh-frozen samples because the limited success of oligo(dT) primers in degraded RNA.^12-16^ Alternatively, 10X Visium Spatial Gene Expression and Patho-DBiT could be applied to FFPE samples but still in a targeted manner or *in situ* RNA polyadenylation.^17^ Spatial methods for host-microbiome interactions, such as SHM-seq and SmT-seq, have been developed by targeted primers to bacterial 16S rRNA or fungal internal transcribed spacer and 18S rRNA.^18,19^ However, these methods still have limitations in capturing whole host and microbial RNAs within the same biological specimen.

Random primers can bind to any location on RNA transcripts, offering an alternative strategy for FFPE and non-polyadenylated microbial samples.^20-23^ Here, we introduce a random primer-based spatial total RNA sequencing method (spRandom-seq), which enables comprehensive profiling of full-length host and microbe total RNAs. With depletion of abundant sequences by hybridization (DASH), we performed CRISPR-based rRNA depletion for mRNA enrichment and dramatically reduced the rRNA percentage.^24,25^ The spRandom-seq captured long non-coding RNAs (lncRNAs), microbial RNAs and other non-polyadenylated transcripts often neglected by conventional workflows and demonstrated uniform distribution across gene bodies. With mouse and clinical FFPE samples, we demonstrated that spRandom-seq can seamlessly integrate with the commercially available 10X Visium and other spatial transcriptomics platforms with various resolutions. We also conducted spatial exploration of RNA biology in clinical FFPE breast cancer sections, elucidating its architectural features and identifying several potential ncRNAs implicated in tumorigenesis. Meanwhile, our analysis revealed spatially structured microbial colonization patterns and localized immune responses, identifying bacterial subpopulations with distinct metabolic adaptations and their correlation with neutrophil-rich regions. This approach facilitates a comprehensive understanding of high-sensitivity transcriptomics and enables full-length total RNA analysis from both eukaryotic and prokaryotic sources within archival FFPE tissues.

## Results

### Overview of the spRandom-seq Method for FFPE Tissues

The main workflow of spRandom-seq is shown in Fig. 1. Initially, 5 μm sections of banked FFPE tissue blocks were cut and subjected to deparaffinization, rehydration, and decrosslinking using standard xylene, alcohol wash, and TE buffer protocols (see methods). Then due to bacterial cell walls, we added the permeabilization with tween and lysozyme to achieve lysis of host cells and bacteria. Subsequently, total RNA was converted into cDNA through multiple annealing of random primers and oligo(dT) primers during *in situ* reverse transcription. The different length of random primers was evaluated using bulk test, and the results indicated no significant difference (Fig. S1). The 3’ hydroxyl terminus of the cDNAs was terminated with poly(dA) tails by terminal transferase (TdT), released through RNA degradation, and bound to poly(dT) primers on a spatially barcoded DNA slide using 10X Genomics CytAssist. The single-strand cDNAs were extended on the expression slide with a specific spatial barcode. Finally, barcoded cDNAs underwent denaturation, amplification, and preparation for paired-end sequencing using next-generation sequencing (NGS). Inevitably, the use of random primers will capture rRNA. To reduce the waste of sequencing reads and optimize the workflow of spRandom-seq, we performed rRNA depletion on two FFPE samples with the CRISPR-based strategy (Fig. S2). The rRNA-depletion treatment reduced the rRNA proportion from 46.1% to 20.7% and 41.3% to 16.4% in two independent sections, respectively. Furthermore, we developed comprehensive approaches to analyze protein-coding and non-coding RNAs and characterize tumor heterogeneity in FFPE specimens. We established a novel analytical framework for host-microbe spatial interaction studies, which integrates cross-species genomic alignment with microbial localization validation.

**Fig. 1:**
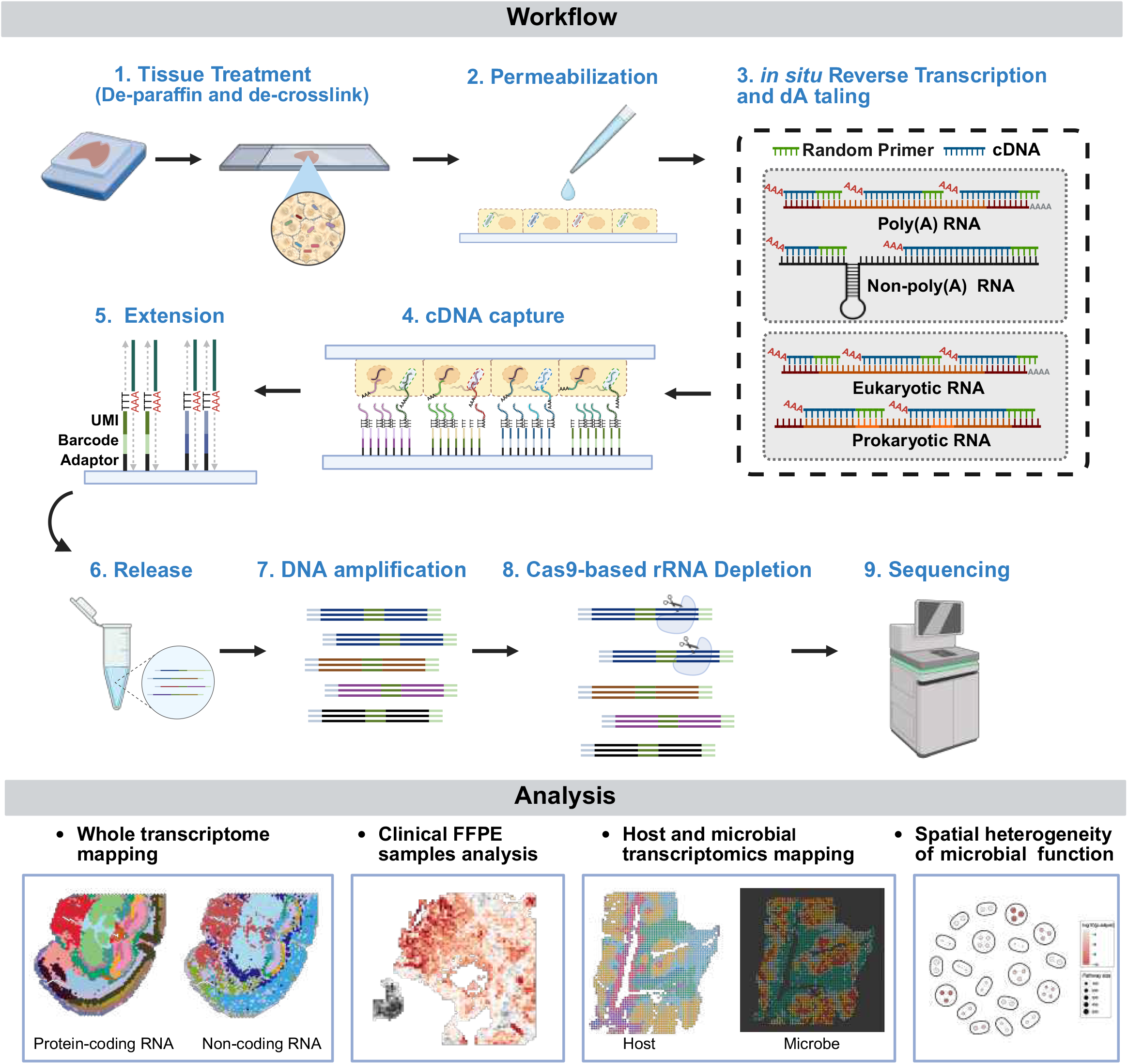
Overview of spRandom-seq workflow for FFPE tissues. The workflow of spRandom-seq for FFPE tissues includes FFPE section pre-treatment, reverse transcription, dA tailing, cDNA transfer, extension, one-strand elution, DNA amplification and sequencing.

### Validation of spRandom-seq Using the Degraded FFPE Mouse Tissues

To evaluate the performance of this technology, we applied spRandom-seq on FFPE sections of mouse coronal brain and heart (the RIN values of these samples were shown in Figure S3) with 10X Visium slides (Resolution: 55 μm). The unsupervised clustering results of our spRandom-seq data aligned perfectly with the annotated mouse brain regions, including the cortex, hippocampus, and striatum, as defined by the Allen Brain Atlas (ABA) using H&E staining (Fig. 2a). We observed comparable results in the mouse heart section, showing the robustness of our methods (Fig. 2a and Fig. S4). At the average sequencing depth of ∼90,000 raw reads per spot, we detected a median of 6,633 UMIs / 2,360 genes and 9,808 UMIs / 2,549 genes for mouse brain and heart, respectively (Fig. 2b). Different cell types were deconvoluted by CARD,^26^ with a validated scRNA-seq dataset from the same tissue as reference (Fig. 2c,d).^27^ The location of each major cell type in spatial location was visualized in Fig. 2c and Fig. S5. For instance, the excitatory neurons specifically expressed the classical cell-type marker gene *Slc17a6* (Fig. 2e). The location of these specific marker genes captured by spRandom-seq was consistent with *in situ* hybridization data from ABA (Fig. 2e).^28^ spRandom-seq further exhibited detection capabilities for various ncRNAs such as *Mir6236* and *Rnu12*, which were inadequately recovered by 10X Visium (Fig. S6a,b). To explore the regional specificity of ncRNAs, we performed unsupervised clustering with sole ncRNAs and the results aligned perfectly with mouse brain regions (Fig. 2f). The major brain regions, such as the isocortex (Cluster 1), dentate gyrus (DG, Cluster 8), CA (Cluster 7 & 9), thalamus (Cluster 2), and hypothalamus (Cluster 3) were well-dissected without protein-coding features, indicating potential region-specific regulatory functions of ncRNAs (Fig. 2f). We performed the differential expression (DE) analysis for each cluster with ncRNAs to further investigate the ncRNAs expressed exclusively in those brain regions (Fig. S6c). Remarkably, *4921539H07Rik* exhibited the highest expression in Cluster 7 (CA1), which recapitulated the previous findings that *4921539H07Rik* is a cell-type specific lncRNA for CA1 excitatory neurons.^29^ Additionally, our data showed that *4921539H07Rik* and *Neurod2* exhibited a similar spatial expression pattern in the CA1 region (Fig. 2g).^29^ These data demonstrated that spRandom-seq offered an approach to spatially characterize the transcriptomic organization of intricate tissues within coding and ncRNAs.

**Fig. 2:**
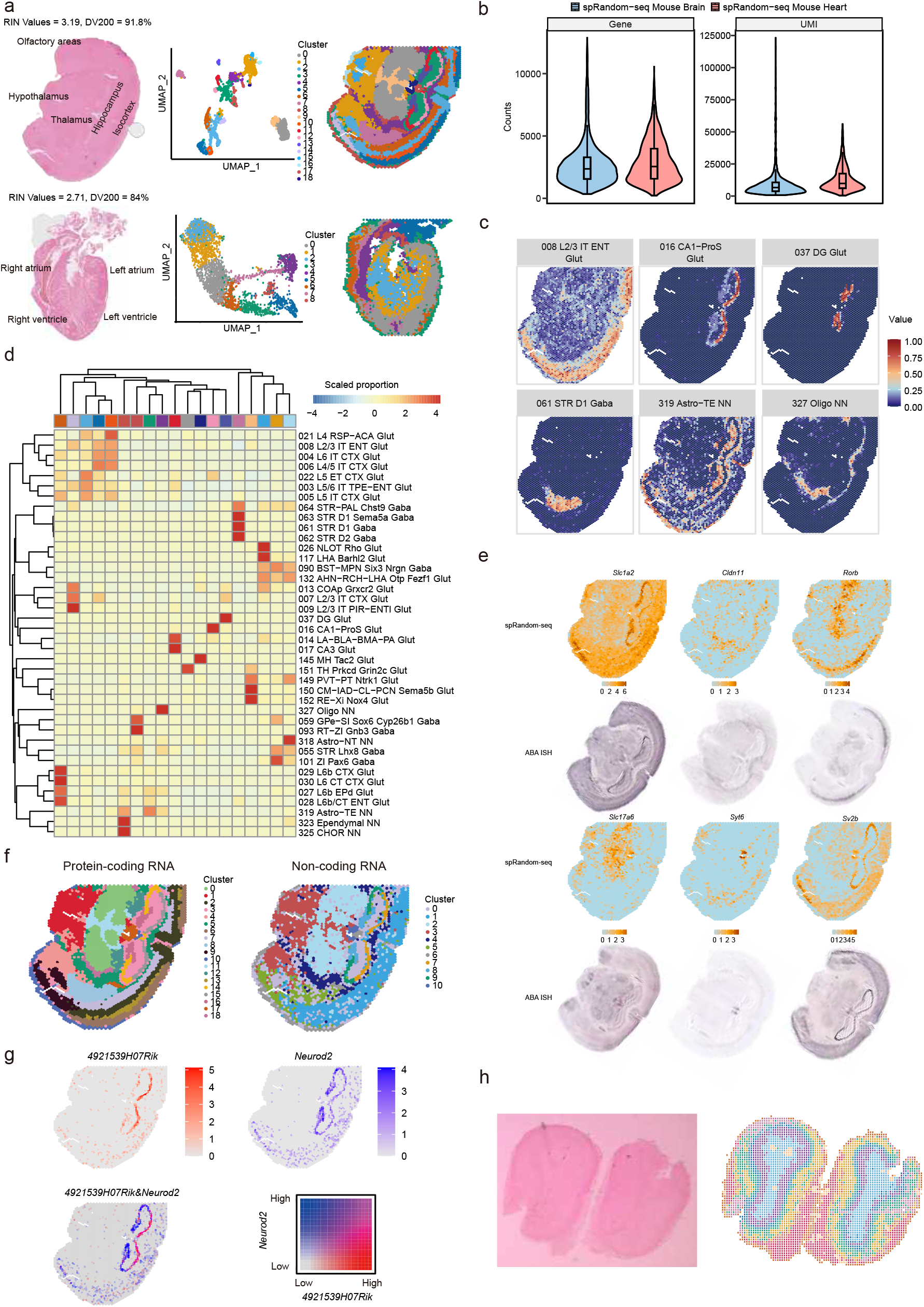
Validation and benchmark of spRandom-seq using a FFPE mouse brain section. **a**, Spatial visualization of coronal FFPE mouse brain and heart section analyzed by spRandom-seq at 55 μm resolution. **b**, Numbers of Genes/UMIs detection for FFPE mouse brain and heart using spRandom-seq. **c**, Spatial maps of classic cell-type for spRandom-seq in FFPE mouse brain section. Spots in which the transcript was not detected are shown as yellow. Color scale indicates log-normalized expression. **d**, Heatmap showing the correlation between clusters and cell types. **e**, Spatial visualization of six selected genes expressions of the FFPE mouse brain section and ISH images of the adult mouse brain taken from ABA. **f**, UMAP of clustering results of coronal FFPE mouse brain section using only protein-coding or non-coding RNAs respectively. **g**, Spatial maps of lncRNA 4921539H07Rik and Neurod2 for spRandom-seq datasets. Spots in which the transcript was identified as lncRNA are depicted in red, while those identified as Neurod2 are depicted in blue. Spots exhibiting co-expression are represented in pink. **h**, Spatial visualization of FFPE mouse olfactory bulb section analyzed by spRandom-seq at 15 μm resolution.

To investigate the performance of spRandom-seq on high-resolution platforms, we applied spRandom-seq on a mouse olfactory bulb FFPE section with a commercially available 15μm-resolution chip and detected a median of 3,700 UMIs and 941 genes. The clustering results successfully aligned with the anatomic structure of mouse olfactory bulb (Fig. 2h). These data indicated spRandom-seq’s high compatibility with various spatial transcriptomics platforms and expression chips with different resolutions.

### Spatial Total Transcriptome Mapping of the Clinical Archival Breast Cancer Using spRandom-seq

spRandom-seq was also employed to elucidate the cellular heterogeneity within tumor tissues obtained from a patient diagnosed with breast cancer. With 44% sequencing saturation, we detected a median of 28,942 UMIs and 5,853 genes per spot for breast cancer section. Unsupervised clustering revealed 9 spatial clusters aligned with histological structures (Fig. 3a). The UMAP delineated distinct cell types defined by canonical markers (Fig. 3b). Our approach exhibited superior performance compared with the 10X Visium fresh frozen sample with similar sequencing depth (Fig. 3c,d and Table S1). spRandom-seq showed no discernible 3’- or 5’-end biases throughout the gene body, in contrast to poly(A)-based 10X fresh-frozen Visium, which displays a clear 3’-end bias when subjected to NGS (Fig. 3e and Fig. S7). These results suggested that our method effectively captured full-length transcripts, highlighting the critical distinction between poly(A)-based and random primer-based strategy.

**Fig. 3:**
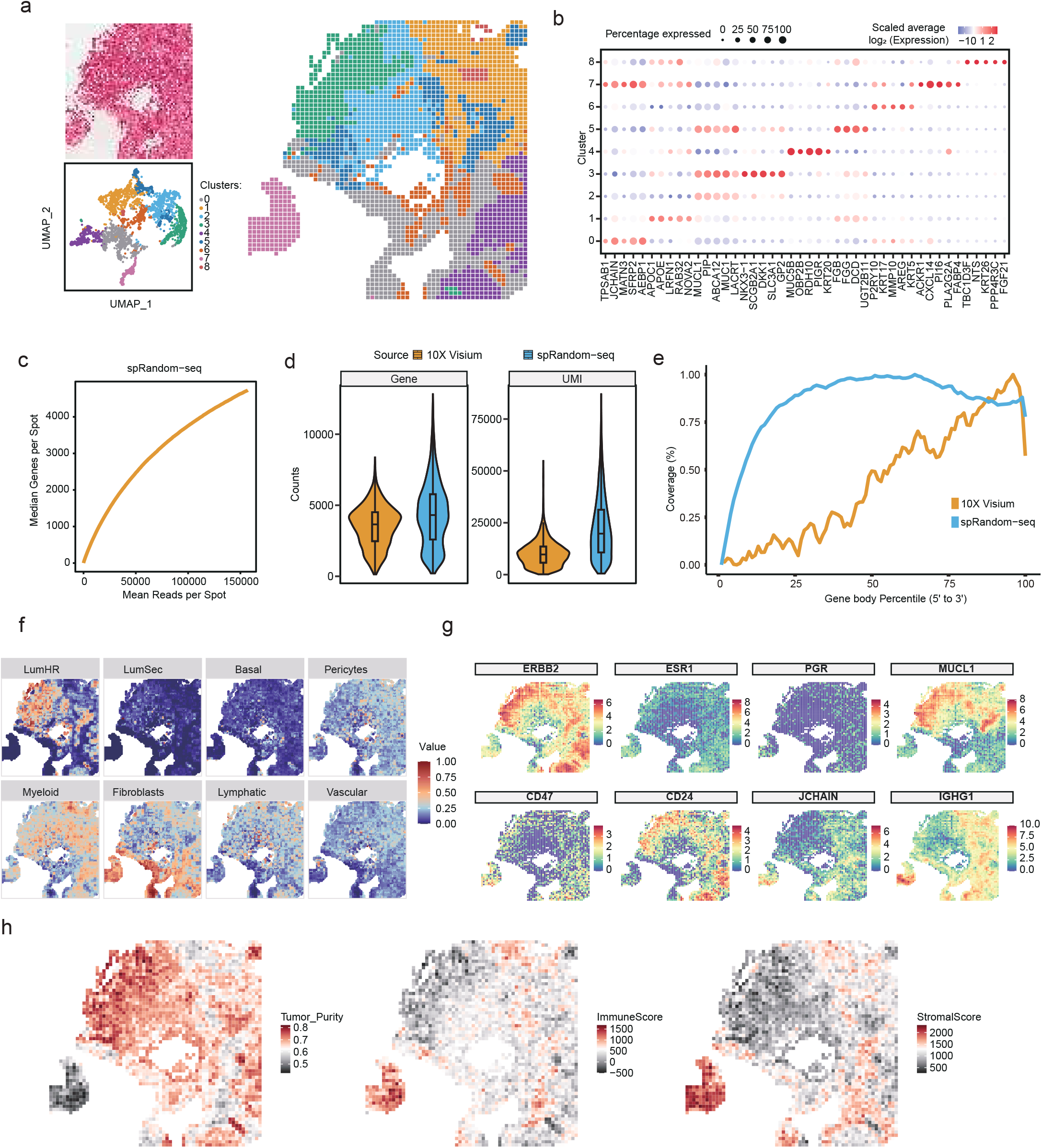
Validation and benchmark of spRandom-seq using a clinical FFPE breast cancer section. **a**, Spatial visualization of the clinical FFPE breast cancer section analyzed by spRandom-seq at 50 μm resolution. **b**, Dotplot of top differentially expressed genes in each cluster based on the unsupervised clustering. **c**, Saturation analysis of spRandom-seq based on the the clinical FFPE breast cancer tissues. **d**, Genes/UMIs detection comparison the clinical FFPE breast cancer tissues using spRandom-seq and fresh frozen breast by 10X Genomics. **e**, Read coverage along the gene body by spRandom-seq in FFPE breast cancer section. **f**, Cell-type deconvolution results displaying spatial distribution of major cell types in breast section. Color scale indicates the proportion of a given cell type in a spot. **g**, Spatial maps of classic cell-type marker genes for spRandom-seq in FFPE breast cancer section. Color scale indicates log-normalized expression. **h**, Spatial maps displaying tumor purity, immune and stromal scores from ESTIMATE analysis of spRandom-seq FFPE breast cancer section.

We performed cell-type deconvolution to obtain the cell-type distribution across the breast cancer section using previous human breast scRNA-seq data.^30^ The location of each major cell type in spatial location was visualized in Fig. 3f. Our data revealed a high expression of *ERBB2* and *ESR1* but low expression of *PGR* in the tumor, suggesting the presence of HER2-positive breast cancer (Fig. 3g). Additionally, the tumor purity score calculated by ESTIMATE accurately identified the specific regions affected by the tumor (Fig. 3h).^31^ We also observed a high expression of *MUCL1* in certain tumor regions, accompanied by low-expression levels of *IGHG1* and *JCHAIN*. Previous reports have suggested that this correlation is associated with worse relapse-free survival in patients.^32^ Additionally, the elevated expression of *CD24* (but not *CD47*) in the tumor area supports its potential as a promising target for cancer immunotherapy in ovarian and breast cancers.^33^ These findings highlight the ability of spRandom-seq to capture the heterogeneity present within clinical FFPE sections of breast cancer.

### Identification of Spatial ncRNAs in the Clinical Archival Breast Cancer Tissues

spRandom-seq exhibited robust detection capabilities for various ncRNAs that were inadequately recovered or undetectable by 10X Visium, including miRNA, snRNA, lncRNA, and miscRNA (Fig. 4a, b). To further elucidate the regulatory role of ncRNAs in tumorigenesis, we first divided the human breast cancer section into “High (Normalized value > 0.5)” and “Low (Normalized value <= 0.5)” regions according to the min-max normalized values of tumor purity, immune score, and stromal score, respectively (Fig. S8a). We then conducted a DE analysis for ncRNAs and protein-coding genes between the high and low tumor/immune/stromal regions (Fig. 4c and Fig. S8b). A Pearson’s correlation test was performed on each pair of significant DE ncRNAs and DE protein-coding genes. Many DE ncRNAs that showed a significant positive correlation with those region-specific protein-coding genes have been reported to be relevant. For instance, the pseudogene *CYP4Z2P* that was highly correlated with DEGs in the high tumor region has been shown to be responsible for maintaining the stemness of breast cancer cells in a previous study (Fig. 4c).^34^ To investigate the potential biological pathways that are associated with those up-regulated ncRNAs in those regions, GO enrichment analysis was performed on the DE protein-coding genes that showed positive correlations with the top upregulated ncRNAs in high tumor/immune/stromal regions respectively (Fig. 4d and Fig. S8c). These were enriched mainly in biological pathway categories associated with epithelial to mesenchymal transition and negative regulation of programmed cell death (Fig. 4e).

**Fig. 4:**
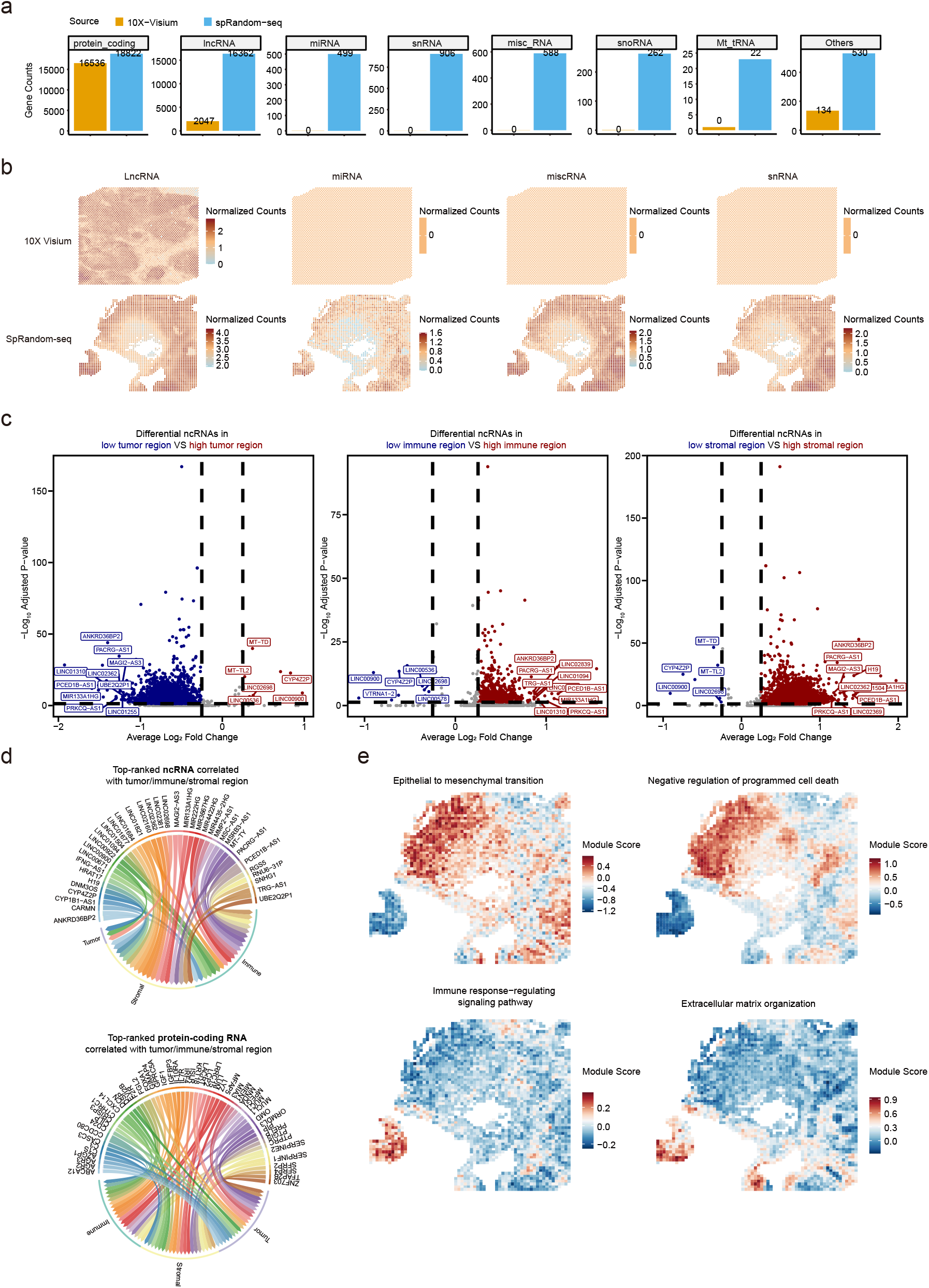
Spatial ncRNA regulation in the clinical FFPE breast cancer section. **a**, Comparison of different RNA biotypes detection between spRandom-seq FFPE breast cancer section and 10X Visium fresh frozen breast cancer section. **b**, Spatial count maps of various ncRNA biotypes. **c**, Volcano plots displaying differentially expressed ncRNAs between the high tumor/immune/stromal and low tumor/immune/stromal regions. **d**, Top-ranked upregulated ncRNAs and protein-coding RNAs showing high correlation with the tumor/immune/stromal regions. **e**, Spatial interactions involving the top upregulated ncRNAs and upstream signaling pathways. The biological pathways were enriched by all protein coding DEGs that showed positive correlation with the ncRNAs. Module score was calculated based on the expression of all intersection genes of a given GO term.

### Validation of spRandom-seq in Capturing Total RNA from Microbes and Hosts

The performance of spRandom-seq was assessed using infected and normal mouse lung FFPE tissues (Resolution: 50 μm). With our single-nucleus RNA-seq (snRandom-seq), unsupervised clustering of the integrated data using classical cell type markers revealed 15 main cell types (Fig. S9a and S9b).^35,36^ The location of each major cell type and gene expressions in spatial location was visualized using CARD (Fig. S10 and 5d).^26^ At an average sequencing depth of ∼30,000 raw reads per spot, we detected a median of 12,046 UMIs/2,997 genes and 10,584 UMIs/2,980 genes in normal and infected mouse lung tissue, respectively. To avoid potential contamination, we set a threshold of 0.05% *K. pneumoniae* (*KP*) transcripts per spot. While *KP* transcripts were undetectable in normal samples, infected samples showed a *KP* distribution map with clear spatial patterns, with some spots containing up to 5% *KP* transcripts (Fig. 5a). At the same sequencing depth, spRandom-seq detected a median of 54 genes and 58 UMI counts for *KP* in the infected sample, with no signal in the normal sample (Fig. 5b). Remarkably, our approach was also effective in detecting *KP* RNA biotypes, recognizing protein-coding RNA, rRNA, transfer messenger RNA (tmRNA), and tRNA (Fig. 5c). We observed that immune cells, especially neutrophils, were significantly enriched in specific areas of the infected samples (Fig. 5d). To confirm that the detected signal specifically indicates *KP* and neutrophils, we conducted IF assays targeting F4/80 for neutrophils and RNA FISH for *KP* (Fig. 5f). The results confirm colocalization and support spRandom-seq’s ability to detect both eukaryotic and prokaryotic RNA. To assess the co-localization of cell types and *KP* distribution, we compared the spatial distribution of all cell types in infected samples and their correlation with *KP* (Fig. 5e). Both capillary 1 cell (CAP1) and alveolar fibroblast 1 (AF1) exhibit a negative correlation with *KP*-infected regions and strongly colocalize (Fig. 5e), indicating their roles in repairing and regenerating infected lung tissue. Notably neutrophils exhibited strong colocalization with the infected lung regions, highlighting their predominant role in the post-infection immune response (Fig. 5g).^37^ In summary, results show the spatial colocalization and correlations between *KP* and host cells in infection, providing insights into the spatial reaction of infection and immune responses.

**Fig. 5:**
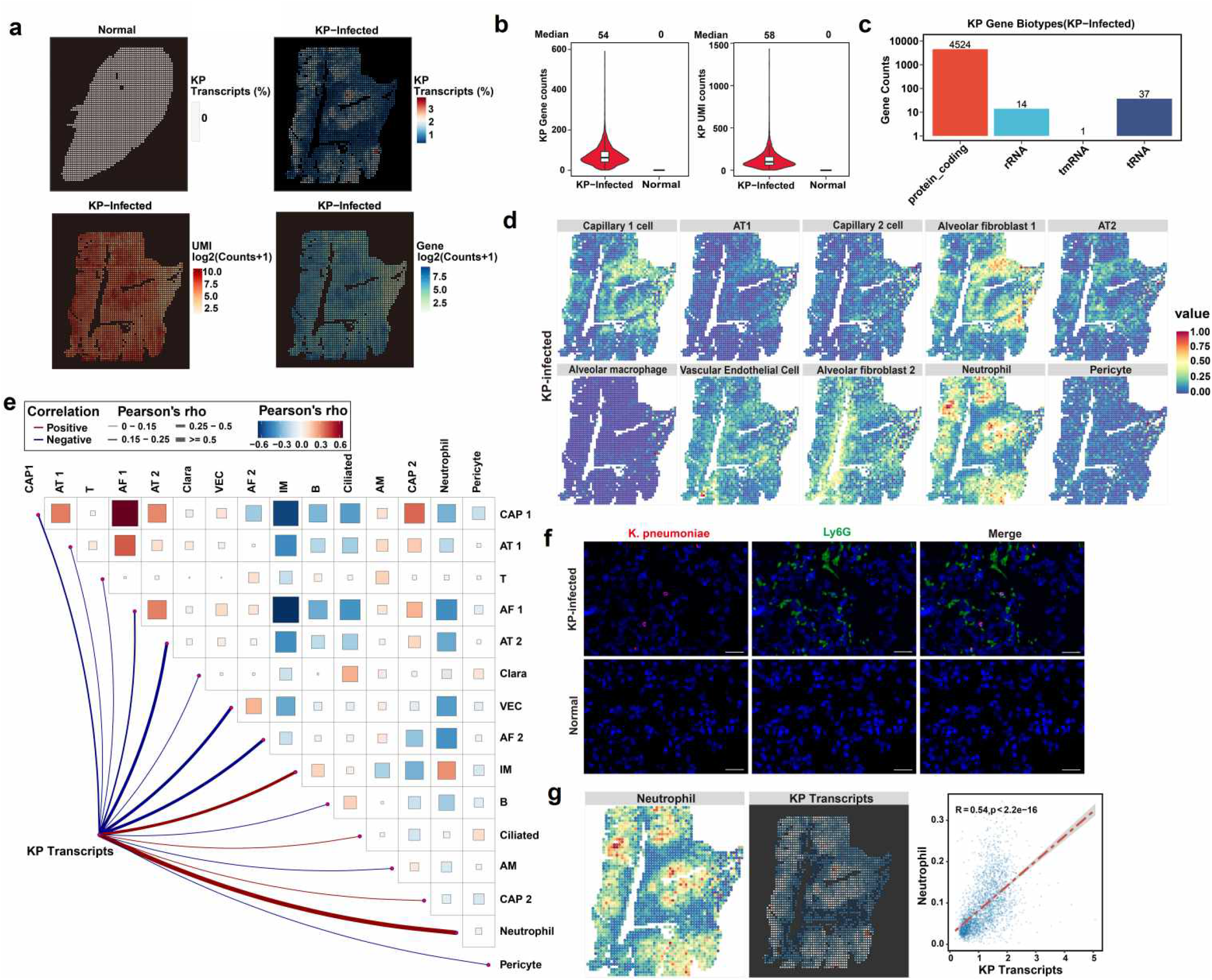
Validation and benchmark of spRandom-seq using normal and infected mouse lungs. **a**, Spatial mappings of the proportion of the KP transcripts in all transcripts per spot in normal and KP-infected samples (Top left and right). Spatial mappings of the Gene counts, and UMI counts of *KP* in KP-infected samples (Bottom right and left). Color scale indicates log-normalized expression. **b**, Numbers of *KP* Genes and UMIs detected per spot in normal and KP-infected samples. **c**, RNA biotypes detected in KP-infected samples. **d**, Cell-type deconvolution results of using snRandom-seq data displaying spatial distribution of major cell types in infected samples. Color scale indicates the normalized proportion of a given cell type in a spot. **e**, Spatial correlation map of the deconvoluted cell-type proportion between cell types (Top right). The correlation analysis between KP transcripts proportion and deconvoluted cell-type proportion of 15 cell types (Bottom left). Correlation is measured by Pearson’s rho. **f**, The lung of normal and infected mouse was analyzed through immunofluorescence and fluorescence *in situ* hybridization. Ly6G (Green), DAPI (Blue), and *KP* (Red). Scale bars, 20 μm. **i**, The Correlation analysis between Neutrophil’s deconvoluted cell-type proportion and *KP* transcripts proportion.

### Spatial Heterogeneity of *K. pneumoniae* Subpopulations Revealed by spRandom-seq

To analyze *KP* heterogeneity, we performed unsupervised clustering and identified three differential subpopulations (Fig. 6a and S11a). Spatial mapping confirmed that clusters 0, 1, and 2 exhibited clear spatial heterogeneity, occupying distinct tissue regions rather than dispersing randomly. The analysis revealed significant variation in microbial load among clusters (Kruskal-Wallis test, p < 2.2e-16), with cluster 0 exhibiting the highest colonization burden, followed by cluster 2 and cluster 1 (Fig. 6b). We performed DE analysis to characterize the transcriptional profile of each cluster, and the top 10 DEGs for each cluster are shown in Fig. S11c. Notably, we found cluster 0 exhibited a higher proportion of immune cell infiltration than clusters 1 and 2 (Fig. S11d). KEGG enrichment analysis showed that cluster 0 had significantly enriched metabolic pathways, including carbon metabolism, nitrogen metabolism, and secondary metabolite biosynthesis. Clusters 1 and 2 had fewer enriched pathways (Fig. 6c). These results implied cluster 0 was experiencing increased environmental pressure, leading to greater enrichment of metabolic pathways.^38-40^

**Fig. 6:**
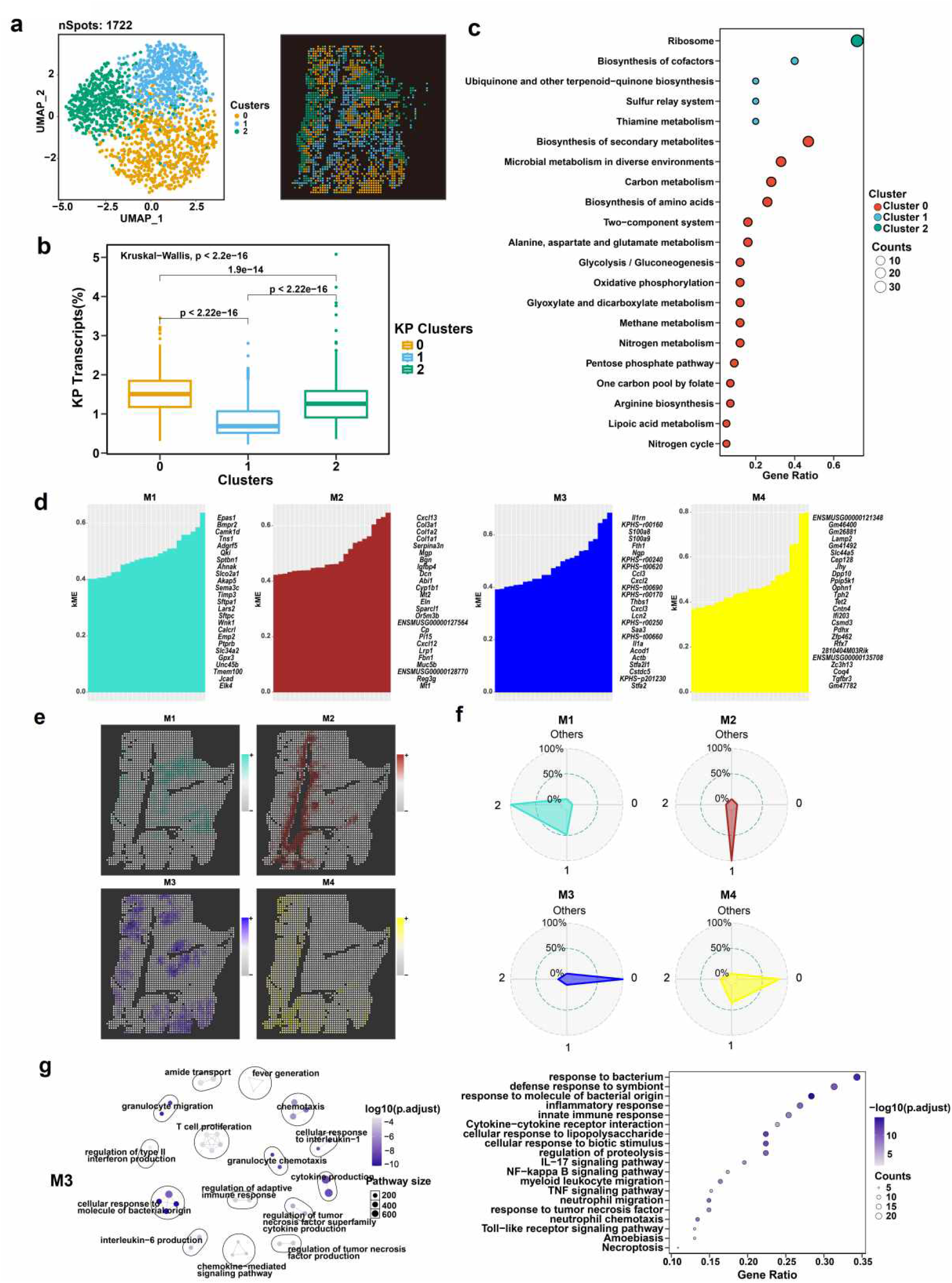
Spatial and functional heterogeneity of *K. pneumoniae* subpopulations in infected samples revealed by spRandom-seq. **a**, UMAP projection and the spatial distribution of major *KP* subpopulations. **b**, The boxplot of KP transcript percentages within clusters 0, 1, and 2. The Kruskal-Wallis test indicates significant differences among clusters (p < 2.2e-16). **c**, Bubble plots of KEGG enrichment analysis for cluster 0,1 and 2 of *KP*. **d**, The hub genes in four modules ranked by kME. **e**, Spatial mappings of module scores calculated by hub genes for the four modules. **f**, The radar plot displaying the relative enrichment of each module in different *KP* clusters. **g**, Top Gene Ontology (GO) terms enriched in the GO Enrichment Analyses with M3 hub genes.

To identify co-expressed gene modules and reveal regional gene expression differences during bacterial infection, we analyzed spRandom-seq data from infected samples using high-dimensional weighted gene co-expression network analysis (hdWGCNA) with a soft threshold power β of 10 (Fig. S12a and S12b).^41,42^ Through dendrogram visualization, we identified a total of 4 distinct gene co-expression modules, including green (M1), red (M2), blue (M3), and yellow (M4) modules (Fig. S12a). We identified hub genes by calculating the eigengene connectivity (kME), and the top 20 genes in each module were visualized (Fig. 6d). In M3, we observed the specific *KP* genes, *KPHS r00160* and *KPHS t00620*, suggesting a potential link between this module and the microbial infection process (Fig. 6d). The *S100a8* and *S100a9* genes were found in M3 and play key roles in bacterial infections, such as antibacterial activity, regulating inflammation, and enhancing immune cell functions. Subsequently, the enrichment fraction of these distinctive hub genes was calculated within spatial maps and indicated that the M3 module was elevated explicitly in the spots where *KP* reads were detected (Fig. 6e). In four infection severity areas, we found that M3 showed significant expression in cluster 0, while M2 had higher expression in cluster 1 (Fig. 6f). We extracted the top 100 significant genes from the four modules for Gene Ontology (GO) enrichment analysis (Fig. 6g and S12c). For M3, hub genes were significantly enriched in pathways related to “response to bacterium,” “defense response to symbiont,” and “response to molecule of bacterial origin,” involving immune and inflammatory processes such as cellular response to bacterial molecules, chemotaxis, and cytokine production (Fig. 6g). In M1, hub genes were primarily associated with vascular development and cell movement, crucial for angiogenesis and differentiation (Fig. S12c). For M2, hub genes were linked to extracellular matrix organization and cell movement, important for migration and tissue development (Fig. S12c). In summary, our hdWGCNA analysis of spRandom-seq data from infected samples identified four gene co-expression modules. Module M3 was potentially linked to *KP* infection and showed elevated expression in cluster 0. The results also indicated that microbial infection induces localized immune responses under spatial regulation.

## Discussion

We developed a spRandom-seq technology by simultaneously capturing full-length total host and microbial RNAs with random primers. This innovative approach offers a potential spatial platform for analyzing full-length total RNA from both eukaryotic and prokaryotic sources within archival FFPE tissues. spRandom-seq leverages widely used spatial gene expression chips such as 10X Visium platform, ensuring ease of operation and commercial scalability for large-scale applications.

Current spatially resolved transcriptomics techniques mainly rely on 3’-end poly(A) tails of mRNA, which are limited to non-degraded and polyadenylated RNA samples. 10X Visium Spatial Gene Expression utilizes probe-based technology to capture spatial information for genes of interest, albeit with limited coverage of the whole transcriptome. In contrast, spRandom-seq enables unbiased and efficient spatial transcription profiling on host and microbial RNA within FFPE samples. Notably, spRandom-seq achieves high sensitivity for microbial transcripts (detecting ∼5% bacterial reads in infected spots), and its integration with snRandom-seq enhances spatial deconvolution, overcoming resolution limitations in spatial transcriptomics. In our study, we identified cellular heterogeneity and microbial subpopulation, revealing a strong spatial correlation between *KP* clusters and neutrophil infiltration. Distinct *KP* subpopulations, enriched in metabolic pathways such as carbon, nitrogen, and glyoxylate metabolism, suggest bacterial adaptation to host microenvironments drives pathogen survival and virulence, aligning with prior studies on metabolic flexibility enhancing resistance to host defenses.

Additionally, the utilization of random primers in spRandom-Seq facilitates enhanced coverage of gene body regions, thereby enabling the identification of a substantial number of ncRNAs and facilitating the exploration of nascent transcripts. The availability of these comprehensive full-length total spatial RNA-seq datasets suggests the potential for a thorough analysis of copy number variation (CNV) and mutations at the spatial level.

Our method has a limitation on the resolution. The resolution of the expression slides in our experiments was about 55 and 15 μm, preventing single-cell analysis for several cell types and microbes. To achieve single-cell resolution, it is imperative to employ expression slides with higher spatial resolution in our methodology.

In conclusion, the straightforward experimental protocol and comprehensive spatial transcriptomic on host and microbial information obtained from the FFPE tissues described in this study are anticipated to facilitate the future large-scale application of spRandom-seq in both basic and clinical research endeavors.

## Methods

### Tissues

Male wildtype C57BL6/J mice (6–8 weeks of age) were ordered from Shanghai SLAC Laboratory Animal. Animals were euthanized with CO_2_ followed by cervical dislocation. Tissues were quickly dissected and soaked in polyformaldehyde (PFA) overnight at 4℃. FFPE samples of mouse tissues were prepared by HaoKe Biotechnology Co. Ltd. The procedures involving mice in this study were approved by the Research Ethics Committee of the First Affiliated Hospital, Zhejiang University School of Medicine (approval numbers: 20231446). The collection of human samples and research conducted in this study was approved by the Research Ethics Committee of the First Affiliated Hospital (approval numbers: IIT20210078B). Informed consent for collection and research using surgically removed adult tissues was obtained from each patient before the operation. All the protocols used in this study were in strict compliance with the legal and ethical regulations of Zhejiang University School of Medicine and Affiliated Hospitals. All the protocols used in this study complied with the “Interim Measures for the Administration of Human Genetic Resources” administered by The Ministry of Science and Technology and The Ministry of Public Health.

### Bacterial strains and culture conditions

Klebsiella pneumoniae AS1.1736, obtained from the China General Microbiological Culture Collection Center (CGMCC; Beijing, China), was used in this study. The KP strain was individually inoculated into 5 mL of fresh Luria-Bertani (LB) liquid medium and incubated at 37°C with shaking at 150 rpm. The optical density at 600 nm (OD600) was measured using a spectrophotometer. When the OD600 reached 0.6-0.8, bacterial cells were harvested for subsequent study.

### Murine Pneumonia Model

All animal experiments in this study were approved by the Research Ethics Committee of the First Affiliated Hospital, Zhejiang University School of Medicine (approval numbers: 20231446). The female C57BL/6 (4 weeks of age) mice were purchased from Lingsci Biotechnology (Kunshan) Co. Ltd. All mice acclimated for a week in a controlled environment with 12 h light per day, air humidity of 40–70%, and temperature between 20°C and 26°C. Following the induction of respiratory anesthesia in mice in day 0, the dose of 10^7 CFU KP suspension in 50ul PBS was administered via intranasal instillation to establish a pneumonia model. This method allows for the controlled introduction of the pathogen and facilitates the study of infection dynamics and host response. In day 3, the mice succumbed to the infection. Mouse lung tissues were harvested under sterile conditions and immediately placed in 4% polyformaldehyde (PFA) in PBS at 4℃ for fixation overnight.

### FFPE sections treatment

FFPE samples of mouse tissues were prepared by HaoKe Biotechnology Co. Ltd. FFPE sections were cut from the paraffin block and were kept on at 60℃ for 2h. The sections were then washed twice with 4 ml Xylene for 20 min at room temperature. The sections were immersing in a graded series of ethanol solutions from pure 100% to 30% for 2 min each time. The sections were washed three times with 4 ml pre-cold wash buffer (1X PBS with 1U/μL RNase inhibitor) at room temperature for 5 min.

The FFPE sections placed on the chip were decrosslinking using pH 9.0 TE buffer, incubated at 95℃ for 60 min and then washed three times with 2X SSC buffer (2X SSC buffer with 1U/μL RNase inhibitor).

The FFPE sections with bacterium were washed and incubated with a solution of 0.04% Tween-20 in PBS containing an RNase inhibitor. The permeabilization of the bacterial cell wall was then performed using 2.5 mg/mL lysozyme. Following the permeabilization step, the sections were promptly washed three times with 2X SSC buffer.

### Reverse transcription

The reverse transcription mix was prepared: 55 μL of ddH_2_O, 20 μL of 5X reverse transcription buffer, 5 μL of 100 mM dNTP, 10 μL of 10 μM random primer, 5μL of Murine RNase Inhibitor, 5μL of reverse transcriptase. The FFPE sections were incubated with the reverse transcription mix with multiple annealing ramping from 8℃ to 42℃ and overnight at 42℃. After incubation, the sections were washed with 2X SSCT (2X SSC with 0.05% T-ween 20) three times to wash away the residual random primers and primer dimers.

### dA tailing

The dA tailing mix was prepared: 78 μL of ddH_2_O, 10 μL of 10X TDT buffer, 10 μL of CoCl_2_ buffer, 1 μL of 100mM dATP, 1 μL of TdT enzyme. The FFPE sections were incubated with the dA tailing mix at 37℃ for 30 min and then were washed with 2X SSCT three times.

### cDNA transfer

The transfer mix was prepared: 34.5 μL ddH_2_O, 5 μL RNase H reaction buffer, 5 μL 1mg/mL Proteinase K, 3 μL RNase H and 2.5μL 10% TritonX 100. Following dA tailing, the cDNA in the FFPE sections were transferred into a gene expression slide (The 50 and 15 μm expression slides were purchased from M20 Genomics) with the transfer mix at 37℃ for 30 min using the 10X CytAssist or M20 Genomics Transfer. The details were following the manual protocols respectively.

### DNA extension in situ

The extension mix was prepared: 72 μL ddH_2_O, 10 μL 10X buffer, 10 μL 10mM dNTP and 8 μL DNA Polymerase. The gene expression slide were incubated with the extension mix at 60℃ for 60 min.

### Library preparation

Following DNA extension, the standard 10X Visium CytAssist library preparation was followed to generate cDNA and final sequencing libraries: The single cDNA containing the spatial barcode was eluted using 50 μl 0.08 M KOH and 3 μl 1 M Tris-HCl pH 8.0. Amplify the mix using 10X Genomics pre-amplification mix for 10 cycles and cleanup by SPRIselect. The cycles was: 98℃ 3 min; 10/rest cycles of 98℃ 15 s, 63℃ 20 s, 72℃ 30 s; End cycles, 72℃ 1 min; 4℃ forever. The rest cycle number of determinations was performed by qPCR and final sequencing libraries was amplified by dual index primers. The libraries were then pooled and sequenced using the NovaSeq X (Illumina).

### Immunofluorescence (IF)-RNA fluorescence in situ hybridization (FISH) double labeling

The Fluorescent in Situ Hybridization Kit was used following the SweAMI probe-FISH□+□IF protocol. The steps were as follows: After deparaffinizing and hydrating the paraffin sections, the sections were boiled in repair solutions for 15□min. Proteinase K (20□μg/ml) was added for 20-30□min. After rinsing with water, PBS washing was performed three times for 5□min each. Prehybridization solution was added and incubated at 37□°C for 1□h. The prehybridization solution was removed, and the hybridization solution containing the probe was added, followed by overnight hybridization at 37□°C. For immunofluorescence staining, sections were blocked with 3% BSA for 30 min, incubated with primary antibody overnight at 4°C, washed with PBS and incubated with fluorescein-labeled secondary antibody for 50 min at room temperature. Nuclei were counterstained with DAPI (RT, 8 min, dark), washed with running water, and mounted with anti-fade medium. Fluorescent microscopy was used to collect images with the following wavelengths: UV (excitation 330–380 nm, emission 420 nm), FAM (488) (excitation 465–495 nm, emission 515–555 nm), and CY5 (excitation 608–648 nm, emission 672–712 nm).

## Computational Methods

### Preprocessing and alignment of the sequencing data for mouse coronal brain and heart samples

To pre-process the mouse coronal brain section and heart data, the barcode (BC) and UMI sequences for each read were extracted from the fastq file for R1 based on their standard lengths (16bp BC + 12bp UMI). For read 2, the polyA sequence was trimmed for each read and we limited the read length (after polyA removal) to be 90bp at most to match the read length of typical 10X Visium results. The processed fastq files were then aligned to the mouse reference genome (mm39, https://ftp.ebi.ac.uk/pub/databases/gencode/Gencode_mouse/release_M32/GRCm39.primary_assembly.genome.fa.gz) downloaded from GENCODE with STARsolo (v2.7.10b). The barcode whitelist file was retrieved from the space ranger software (v2.1.0, visium-v4_coordinates.txt).

### Preprocessing and alignment of the sequencing data for clinical human breast cancer ffpe samples

To pre-process the clinical human breast cancer FFPE data, the barcode (BC) and UMI sequences for each read were extracted from the fastq file for R1 based on their lengths (20bp BC + 8bp UMI). For read 2, the adapter and polyA sequences were trimmed for each read. The processed fastq files were then aligned to the human reference genome (hg38, https://ftp.ebi.ac.uk/pub/databases/gencode/Gencode_human/release_43/GRCh38.primary_assembly.genome.fa.gz) downloaded from GENCODE with STARsolo (v2.7.10b).

### Preprocessing and alignment of the msprandom-seq sequencing data for the murine pneumonia model

To pre-process the sequencing data of murine model of KP-induced pneumonia, the barcode (BC) and UMI sequences for each read were extracted from the fastq file for R1 based on their lengths (20bp BC + 8bp UMI). For read 2, the adapter and polyA sequences were trimmed for each read. The processed fastq files were then aligned to the integrated reference genome of mouse and KP with STARsolo (v2.7.10b). The mouse genome (mm39, https://ftp.ebi.ac.uk/pub/databases/gencode/Gencode_mouse/release_M32/GRCm39.primary_assembly.genome.fa.gz) was downloaded from GENCODE. The reference genome of KP was downloaded from RefSeq (https://www.ncbi.nlm.nih.gov/datasets/genome/GCF_000240185.1/).

### Spatial transcriptomics sequencing data from other technologies

For mouse sample, the sequencing data for 10X Visium FF mouse coronal brain section data was downloaded from the 10x Genomics website (https://www.10xgenomics.com/resources/datasets/adult-mouse-brain-coronal-section-fresh-frozen-1-standard) and was processed with the same pipeline that we used for spRandom-seq. The mm39 genome annotation file downloaded from GENCODE (https://ftp.ebi.ac.uk/pub/databases/gencode/Gencode_mouse/release_M32/gencode.vM32.primary_assembly.annotation.gtf.gz) was used to annotate gene biotypes. For human breast cancer sample, the filtered expression matrix for 10X Fresh/Frozen human breast cancer section data was retrieved from 10X Genomics website (https://www.10xgenomics.com/datasets/human-breast-cancer-visium-fresh-frozen-whole-transcriptome-1-standard). According to the provided summary file, same human genome (GRCh38) was used to perform the sequence alignment. The hg38 genome annotation file downloaded from GENCODE (https://ftp.ebi.ac.uk/pub/databases/gencode/Gencode_human/release_43/gencode.v43.primary_assembly.annotation.gtf.gz) was used to annotate the gene biotypes.

### Data processing of the snrandom-seq murine pneumonia model data

For all data generated by snRandom-seq, the genes expressed in fewer than 3 cells, and the cells with fewer than 200 genes detected were removed. To further exclude cells that were highly contaminated by ambient RNAs, we calculated contamination rate for each cell with decontX (v1.0.0) packages in R. The cells with contamination rate higher than 0.2 were removed from the expression matrix. For all the data, the identification of top 3000 variable features (‘FindVariableFeatures’), the normalization (‘SCTransform’), dimension reduction (‘runPCA’, ‘runUMAP’), the shared nearest-neighbor (SNN) graph construction using (‘FindNeighbors(dim = 1:30)’), and Louvain clustering (‘FindCluster’, algorithm = 1) were performed with built-in functions in Seurat (v5.1.0) package in R (v4.3.3). To perform the differential expression (DE) analysis with ‘FindAllMarkers(only.pos = TRUE, min.pct = 0.1, logfc.threshold = 0.25)‘ function in Seurat, scaling and normalization were re-performed on the RNA assay with ‘NormalizeData()‘ and ‘ScaleData()‘ function in Seurat. Only the DE genes that passed 0.05 threshold after false discovery rate correction were retained for downstream analyses.

The visualization of scRNA-seq data was done by the SCP (v0.5.6) package in R.

### Data processing of the sprandom-seq sequencing data

For all data generated by spRandom-seq, the spots under tissue were first selected manually based on the H&E images. For all the data, the identification of top 3000 variable features (‘FindVariableFeatures’), the normalization (‘SCTransform’), dimension reduction (‘runPCA’, ‘runUMAP’), the shared nearest-neighbor (SNN) graph construction using (‘FindNeighbors(dim = 1:30)’), and Louvain clustering (‘FindCluster’, algorithm = 1) were performed with built-in functions in Seurat (v5.1.0) package in R (v4.3.3). To perform the differential expression (DE) analysis with ‘FindAllMarkers(only.pos = TRUE, min.pct = 0.1, logfc.threshold = 0.25)‘ function in Seurat, scaling and normalization were re-performed on the RNA assay with ‘NormalizeData()‘ and ‘ScaleData()‘ function in Seurat. Only the DE genes that passed 0.05 threshold after false discovery rate correction were retained for downstream analyses.

### Cell deconvolution of the data generated by sprandom-seq

The cell deconvolution of the spRandom-seq data were done by CARD package (v1.1) in R with all default parameters. The reference single-cell RNA-seq data for mouse brain was the Adolescent Mouse Brain dataset downloaded from the Mouse Brain Atlas (https://mousebrain.org/). The cell type annotation of Mouse Brain Atlas dataset was reannotated with the MapMyCells tool (https://portal.brain-map.org/atlases-and-data/bkp/mapmycells) developed by Allen Institute with Allen Brain Cell Atlas datasets to obtain more detailed cell-type annotation.[10] The reference single-cell RNA-seq data for mouse heart was downloaded from Tabula Muris (https://tabula-muris.ds.czbiohub.org/). For human breast cancer sample, we used the human breast single-cell RNA-seq dataset generated by Kumar et al., and the dataset was downloaded from the CELLxGENE database with the following link (https://cellxgene.cziscience.com/collections/4195ab4c-20bd-4cd3-8b3d-65601277e731).^30^ For the data of murine pneumonia model, the corresponding snRandom-seq data was used as reference data.

The cell type proportion plot was generated with the ‘CARD.visualise.prop‘ function in CARD package.

### Genebody coverage

The gene body coverage from 5’ end to 3’ end for each sample was calculated by using the ‘genebody_coverage.py’ script from RSeQC (v4.0.0) software.

### Calculation of tumor purity for clinical sample with ESTIMATE

To calculate the tumor purity for the human breast cancer data, we performed the ESTIMATE analysis with the estimate (v1.0.13) package in R. ESTIMATE is a software that calculates the tumor purity, the infiltration level of immune cells in tumor tissue, and the abundance of stromal cells based on expression data. Based on the gene signatures derived from immune and stromal cells, ESTIMATE analysis provides three scores: ESTIMATE score, immune score, and stromal score. The ultimate tumor purity can be calculated with the formula (‘Tumour purity=cos (0.6049872018+0.0001467884 × ESTIMATE score)’) provided in Yoshihara et al.^31^

To further classify regions on the human breast section for downstream analysis, we applied min-max normalization on the tumor purity, immune score and stromal score to ensure the normalized scores are in the range from 0 to 1. Then we classify the spots on the section into High (normalized score > 0.5) and Low (normalized score <= 0.5) region according to the normalized tumor purity, immune score and stromal score respectively.

### Microrna alignment and quantification

The human microRNA annotation file was downloaded from the miRBase database with the following link (https://www.mirbase.org/download/hsa.gff3).[20] We then searched through the STAR-aligned reads in the BAM file to find reads that overlapped with microRNAs. The duplicated reads that share the same barcode and UMI sequences were removed. The filtered BAM was then used to quantify the expression level of microRNAs.

### Differential expression analysis on protein coding genes and non-coding rnas

In order to identify differentially expressed genes and ncRNAs within the pre-defined regions (“Tumor High”, “Immune High”, “Stromal High”), we performed DE analysis using the ‘FindAllMarkers‘ function in Seurat package without any p-values and log2 fold change cutoffs at the first place. The significant DE genes and ncRNAs were then selected with the standard: adjusted P-value <= 0.05 and averaged log2 fold change >= 0.25. Here, only up-regulated genes and ncRNAs were kept for downstream analyses. The volcano plots were plotted with ggplot2 package in R.

### Pearson’s correlation test between differentially expressed protein coding genes and ncRNAs

For the DEGs identified in each pre-defined region, we selected top50 protein coding genes with highest log2 fold change and most significant p-values. Then we performed the pearson’s correlation test between each pair of the top50 protein coding genes and DE ncRNAs. The ggsankey (v0.0.99999) package in R was used to draw the sankey plot to display correlated ncRNA-protein coding gene pairs within “Tumor High”, “Immune High”, and “Stromal High” regions.

### Gene ontology (GO) enrichment analysis on differentially expressed protein coding genes

For the DEGs identified in each pre-defined region, we kept all protein coding genes for the GO enrichment analysis. The ‘gost‘ function in gProfiler2 (v0.2.3) package was applied for the analysis. We limited the source to biological processes in Gene Ontology database only. Then the top GO terms for each region were selected and the overlapped genes were extracted to calculate the module scores for each selected GO terms based on the expression data of the human breast cancer sample. The module score was calculated by the ‘addModuleScore‘ function in Seurat.

### Spatial and functional heterogeneity of K. Pneumoniae subpopulations in infected samples

To explore the potential spatial and functional sub-populations of KP within the infected samples, we first subset the original expression matrix with solely KP genes. We then performed the identification of top 500 variable features (‘FindVariableFeatures’), the normalization (‘SCTransform’), dimension reduction (‘runPCA’, ‘runUMAP’), the shared nearest-neighbor (SNN) graph construction using (‘FindNeighbors(dim = 1:10)’), and Louvain clustering (‘FindCluster’, algorithm = 1) with built-in functions in Seurat (v5.1.0) package in R (v4.3.3). To perform the differential expression (DE) analysis with ‘FindAllMarkers(only.pos = TRUE, min.pct = 0.05, logfc.threshold = 0.05)‘ function in Seurat, scaling and normalization were re-performed on the RNA assay with ‘NormalizeData()‘ and ‘ScaleData()‘ function in Seurat. Only the DE genes that passed 0.1 threshold after false discovery rate correction were retained for downstream analyses. The cluster 0,1, and 2 have differentially expressed genes and much more specific spatial distribution so they were kept for downstream analysis.

To test the difference in KP infection among cluster 0,1, and 2, we performed the Kruskal-Wallis test and the post hoc pairwise wilcoxon tests with the ‘stat_compare_means‘ function in the ggpubr (v0.6.0) package in R, while plotting the boxplot with the ‘ggboxplot‘ function.

To investigate the functional heterogeneity among cluster 0,1, and 2, we applied the ‘enrichKEGG()‘ function in the clusterProfiler (v4.8.3) package in R to perform the gene ontology enrichment analysis with the KEGG database. The organism code corresponds to the Klebsiella pneumoniae subsp. pneumoniae HS11286 is “kpm”. The query gene set for each cluster was the DE genes passed the filtering criteria mentioned above. The KEGG terms with FDR-corrected p-value less than 0.15 were determined as significantly enriched terms for a certain cluster.

### High-dimensional weighted gene co-expression network analysis on infected samples

To identify co-expressed gene modules and reveal regional gene expression differences and interactions during bacterial infection, we analyzed mspRandom-seq data from infected samples using high-dimensional weighted gene co-expression network analysis (hdWGCNA) with the hdWGCNA (v0.4.00) package in R. When we constructed the meta spot with ‘MetacellsByGroups()‘ function, we used the spatial coordinates as the dimensionality reduction result, set the number of the nearest neighbors to be 10 (k=10), and allowed maximum of 6 (max_shared = 6) shared spots between two meta spots. We then followed the basic workflow of hdWGCNA to normalize the data, find the optimized power, and identify hub genes within significant modules based on their intra-module connectivity.

### Gene ontology analysis of module hub genes

To determine the underlying biological functions of the co-expressed gene modules identified by hdWGCNA, we performed the gene ontology enrichment analysis with the top 100 hub genes. The ‘gost‘ function in gProfiler2 (v0.2.3) package was applied for the analysis. We limited the source to GO (GO:BP, GO:MF), KEGG, REAC, and CORUM. The GO terms had adjusted p-values less than 0.05 were determined to be significant. The top and most biologically relevant GO terms were selected manually to be visualized and used to generate the enrichment network. The ‘enrichmentNetwork()‘ function in the aPEAR (v1.0.0) package in R was used to generate the enrichment network for each module, while setting clustering method to ‘hierarchical clustering’ and similarity method to ‘cosine similarity’.

## Reporting summary

Further information on research design is available in the Nature Portfolio Reporting Summary linked to this article.

## Data availability

The Visium mouse coronal brain section data can be downloaded from 10x Genomics website (https://www.10xgenomics.com/resources/datasets/adult-mouse-brain-coronal-section-fresh-frozen-1-standard). The 10X Visium Fresh/Frozen human breast cancer section data can be retrieved from 10X Genomics website (https://www.10xgenomics.com/datasets/human-breast-cancer-visium-fresh-frozen-whole-transcriptome-1-standard). The spatial total transcriptome data generated with spRandom-seq in this study can be found on Gene Expression Omnibus (GEO) and is available under the accession number “GSE247881” (https://www.ncbi.nlm.nih.gov/geo/query/acc.cgi?&acc=GSE247881).

## Code availability

The scripts for data pre-processing and downstream analysis can be found at https://github.com/WangycLab/spRandom-seq.

## Acknowledgements

The project was supported by “Pioneer” R&D programs of Zhejiang Province (No. 2024C03005, Y.W.), the National Natural Science Foundation of China (No. 32200073, Y.W., 32250710678, Y.W. and No. 82200977, Z.X.), Leading Innovative and Entrepreneur Team Introduction Program of Zhejiang (No. 2021R01012, Y.W.), and Key R&D Program of Zhejiang (No. 2024SSYS0022, Y.W.). Thanks for the technical support from the core facilities of Zhejiang University and Liangzhu Laboratory.

## Author Contributions

All authors contributed to the manuscript. Y.L. and J.C. conceived the study. Y.L., S.Z., Y.C., Y.L., B.Z., H.C. and Y.Z. conducted the experiments. Z.X., Y.Y., X.X., N.L. and J.C. analyzed the data and wrote the paper. Y.W., H.G. and J.C. supervised this project. All authors participated in manuscript revision and approved the final manuscript.

