## Supplementary Figures for "Spatial Total RNA Sequencing of Formalin-Fixed Paraffin Embedded Tissues by spRandom-seq"

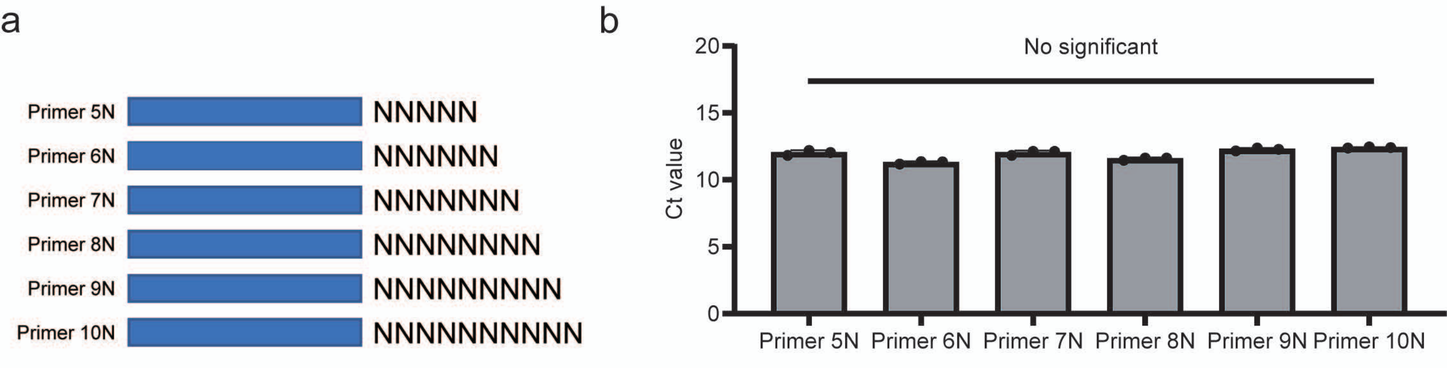
a
Primer 5N
NNNNN
Primer 6N
NNNNNN
Primer 7N
NNNNNNN
Primer 8N
NNNNNNNN
Primer 9N
NNNNNNNNN
Primer 10N
NNNNNNNNNN
b
No significant
Ct value
0
5
10
15
20
Primer 5N
Primer 6N
Primer 7N
Primer 8N
Primer 9N
Primer 10N

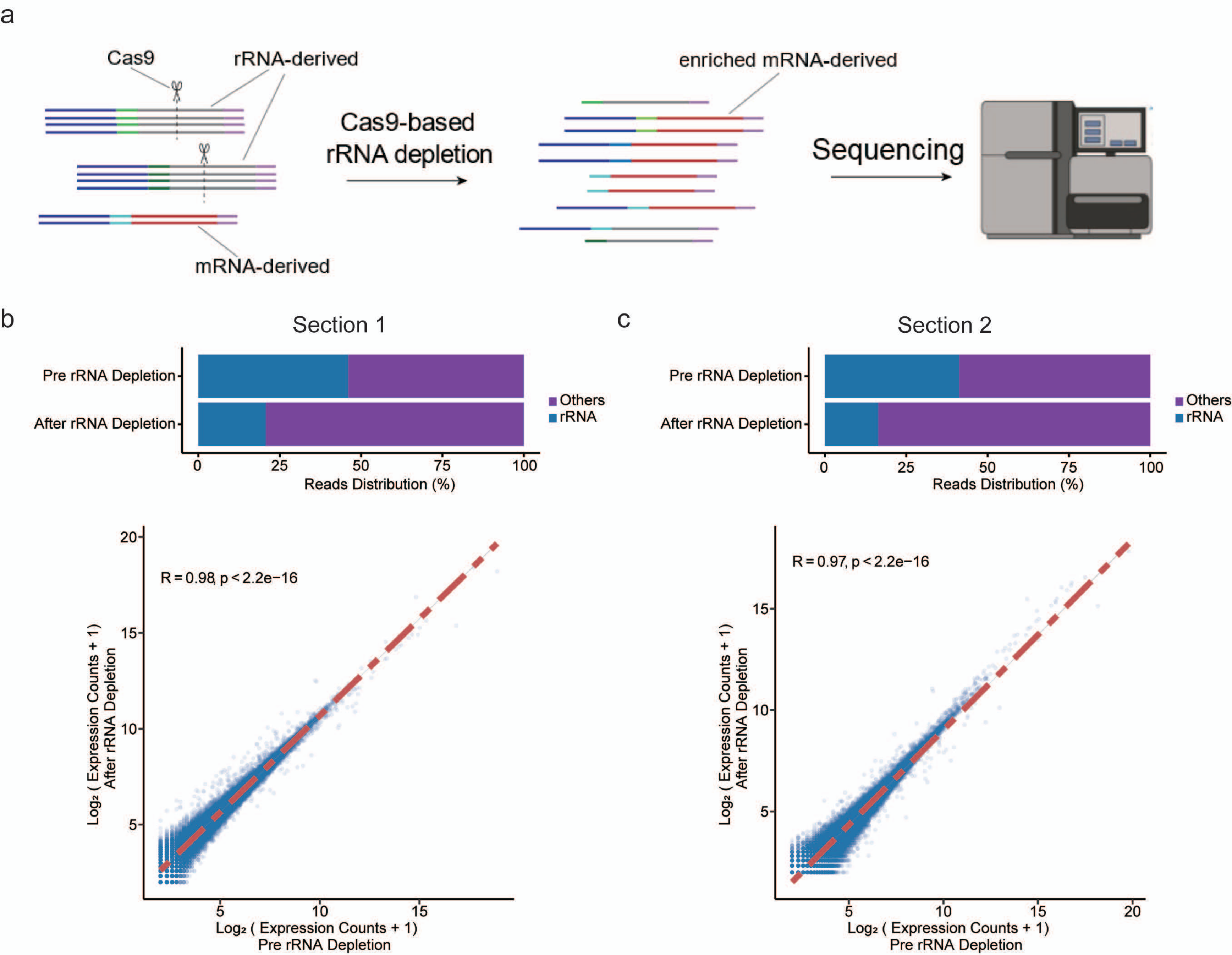
a
Cas9
rRNA-derived
mRNA-derived
Cas9-based rRNA depletion
enriched mRNA-derived
Sequencing
b
Section 1
Pre rRNA Depletion
After rRNA Depletion
Others
rRNA
0 25 50 75 100
Reads Distribution (%)
R = 0.98, p < 2.2e−16
Log₂ ( Expression Counts + 1) After rRNA Depletion
Log₂ ( Expression Counts + 1) Pre rRNA Depletion
c
Section 2
Pre rRNA Depletion
After rRNA Depletion
Others
rRNA
0 25 50 75 100
Reads Distribution (%)
R = 0.97, p < 2.2e−16
Log₂ ( Expression Counts + 1) After rRNA Depletion
Log₂ ( Expression Counts + 1) Pre rRNA Depletion

a

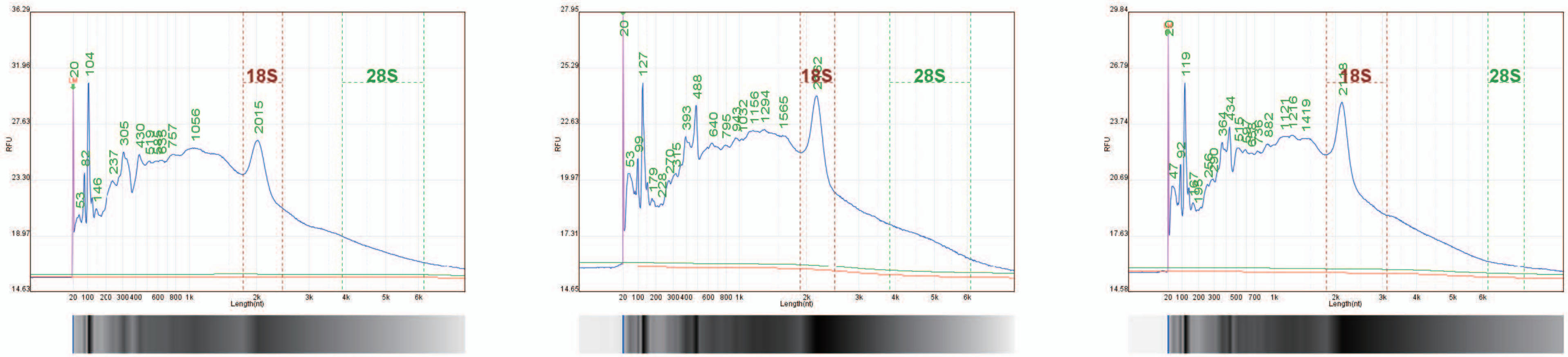


| | Mouse FFPE brain-replicates 1 | | Mouse FFPE brain-replicates 2 | | Mouse FFPE brain-replicates 3 |
|---|---|---|---|---|---|
| Total Peak Area | 31,829,787 | Total Peak Area | 20,635,971 | Total Peak Area | 23,627,412 |
| 18S Area | 4,573,021 (14.4 %) | 18S Area | 2,846,262 (13.8 %) | 18S Area | 4,980,387 (21.1 %) |
| 28S Area | 2,415,382 (7.6 %) | 28S Area | 1,909,053 (9.3 %) | 28S Area | 0.08 |
| Ratio (28S/18S) | 0.53 | Ratio(28S/18S) | 0.67 | Ratio(28S/18S) | 401,817 (1.7 %) |
| RNA Quality Number | 3.14 | RNA Quality Number | 3.23 | RNA Quality Number | 3.21 |
| DV200 | 91.8 % | DV200 | 91.7 % | DV200 | 91.8 % |

b

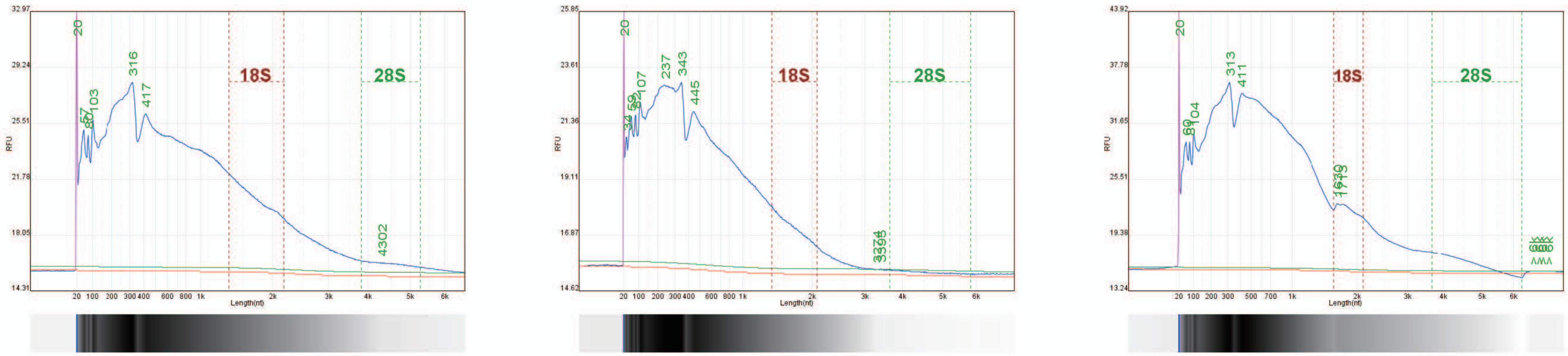


| | Mouse FFPE heart-replicates 1 | | Mouse FFPE heart-replicates 2 | | Mouse FFPE heart-replicates 3 |
|---|---|---|---|---|---|
| Total Peak Area | 24,944,234 | Total Peak Area | 12,721,634 | Total Peak Area | 41,084,390 |
| 18S Area | 3,641,128 (14.6 %) | 18S Area | 1,168,441 (9.2 %) | 18S Area | 2,864,408 (7.0 %) |
| 28S Area | 632,238 (2.5 %) | 28S Area | 129,513 (1.0 %) | 28S Area | 0.44 |
| Ratio(28S/18S) | 0.17 | Ratio(28S/18S) | 0.11 | Ratio(28S/18S) | 1,273,658 (3.1 %) |
| RNA Quality Number | 3.03 | RNA Quality Number | 2.00 | RNA Quality Number | 3.07 |
| DV200 | 84.7 % | DV200 | 79.0 % | DV200 | 85.6 % |

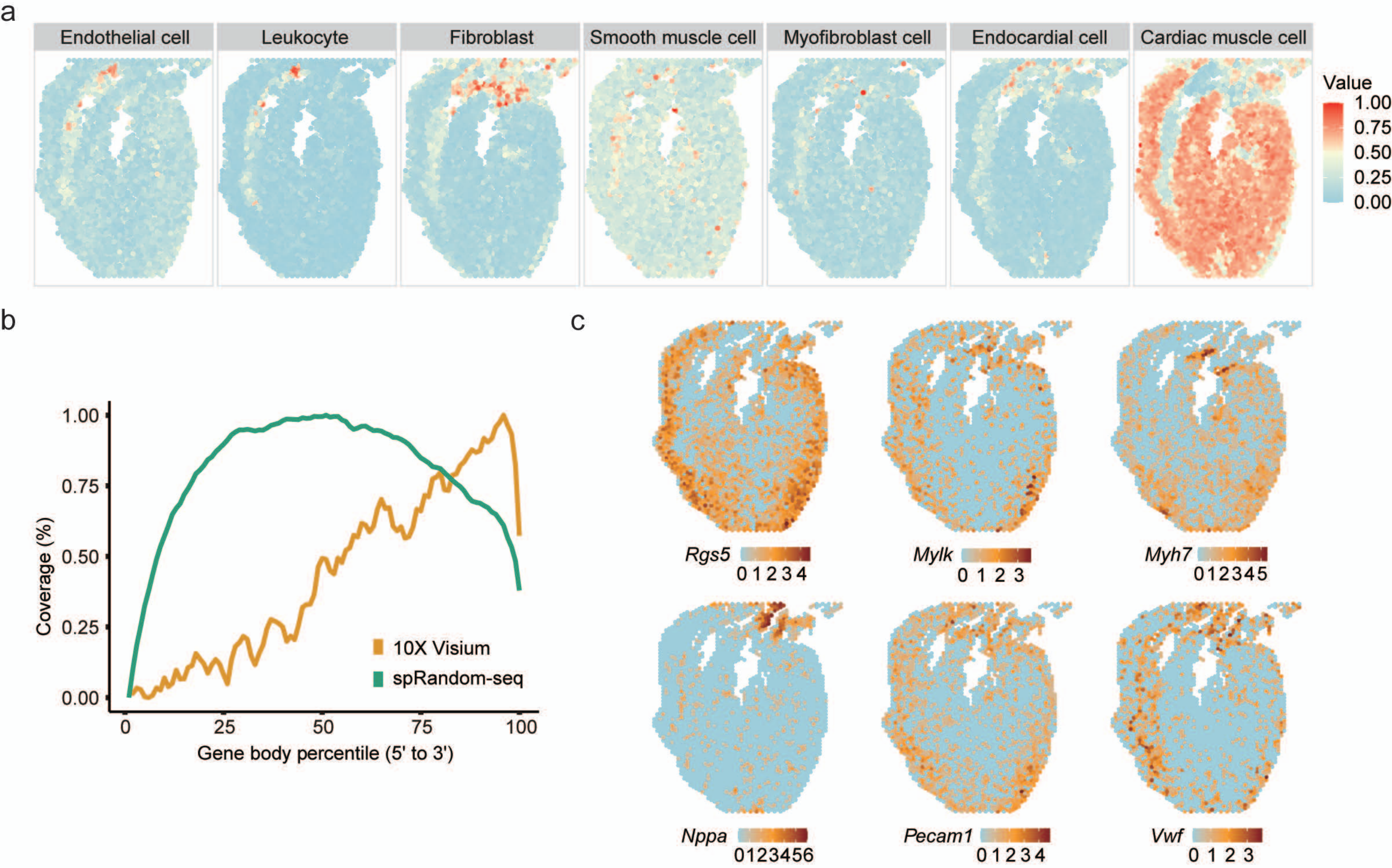
a
Endothelial cell
Leukocyte
Fibroblast
Smooth muscle cell
Myofibroblast cell
Endocardial cell
Cardiac muscle cell
Value
1.00
0.75
0.50
0.25
0.00
b
Coverage (%)
1.00
0.75
0.50
0.25
0.00
0
25
50
75
100
Gene body percentile (5' to 3')
10X Visium
spRandom-seq
c
Rgs5
0 1 2 3 4
Mylk
0 1 2 3
Myh7
0 1 2 3 4 5
Nppa
0 1 2 3 4 5 6
Pecam1
0 1 2 3 4
Vwf
0 1 2 3

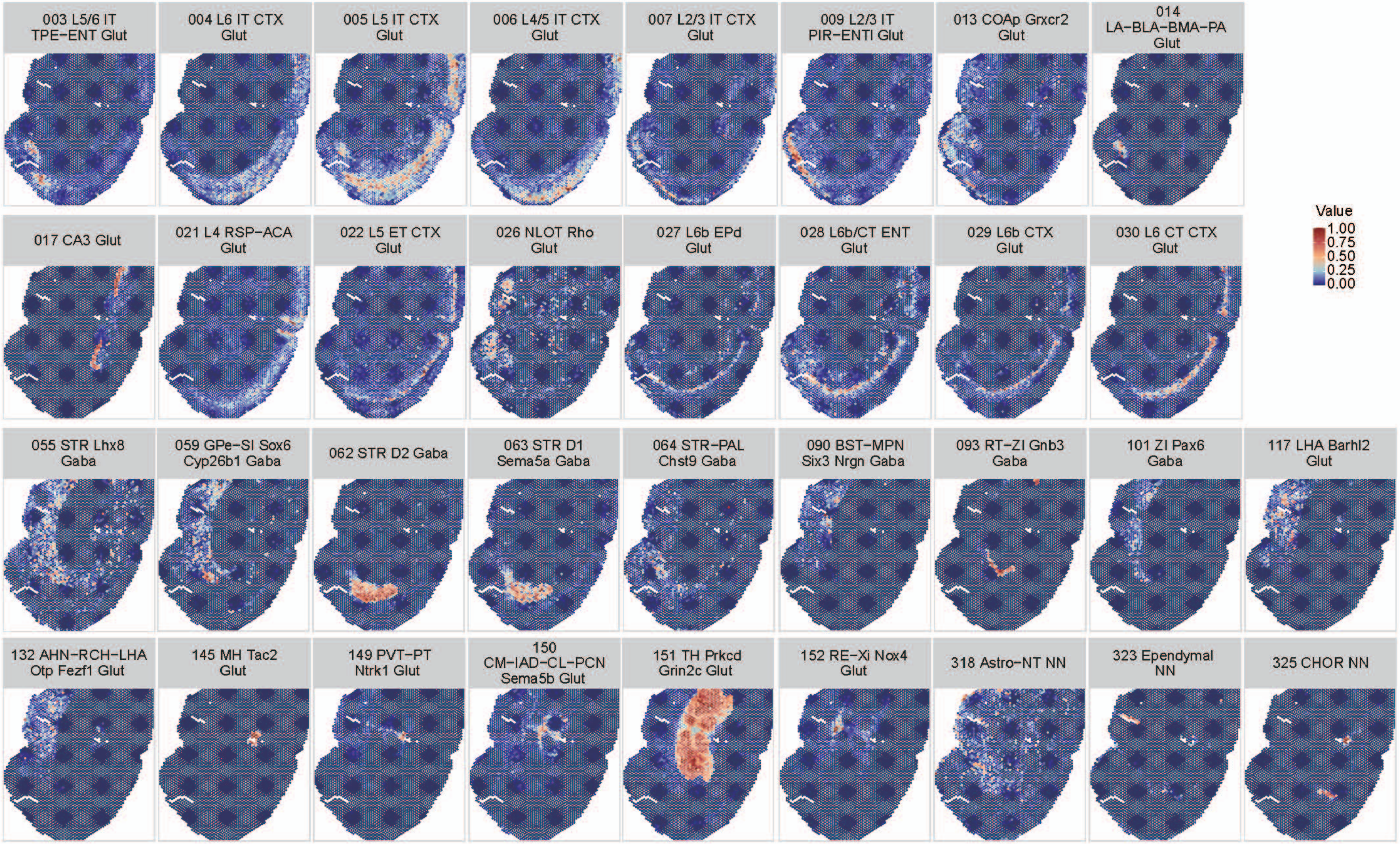

003 L5/6 IT TPE−ENT Glut
004 L6 IT CTX Glut
005 L5 IT CTX Glut
006 L4/5 IT CTX Glut
007 L2/3 IT CTX Glut
009 L2/3 IT PIR−ENTl Glut
013 COAp Grxcr2 Glut
014 LA−BLA−BMA−PA Glut
017 CA3 Glut
021 L4 RSP−ACA Glut
022 L5 ET CTX Glut
026 NLOT Rho Glut
027 L6b EPd Glut
028 L6b/CT ENT Glut
029 L6b CTX Glut
030 L6 CT CTX Glut
055 STR Lhx8 Gaba
059 GPe−SI Sox6 Cyp26b1 Gaba
062 STR D2 Gaba
063 STR D1 Sema5a Gaba
064 STR−PAL Chst9 Gaba
090 BST−MPN Six3 Nrgn Gaba
093 RT−ZI Gnb3 Gaba
101 ZI Pax6 Gaba
117 LHA Barhl2 Glut
132 AHN−RCH−LHA Otp Fezf1 Glut
145 MH Tac2 Glut
149 PVT−PT Ntrk1 Glut
150 CM−IAD−CL−PCN Sema5b Glut
151 TH Prkcd Grin2c Glut
152 RE−Xi Nox4 Glut
318 Astro−NT NN
323 Ependymal NN
325 CHOR NN
Value
1.00
0.75
0.50
0.25
0.00

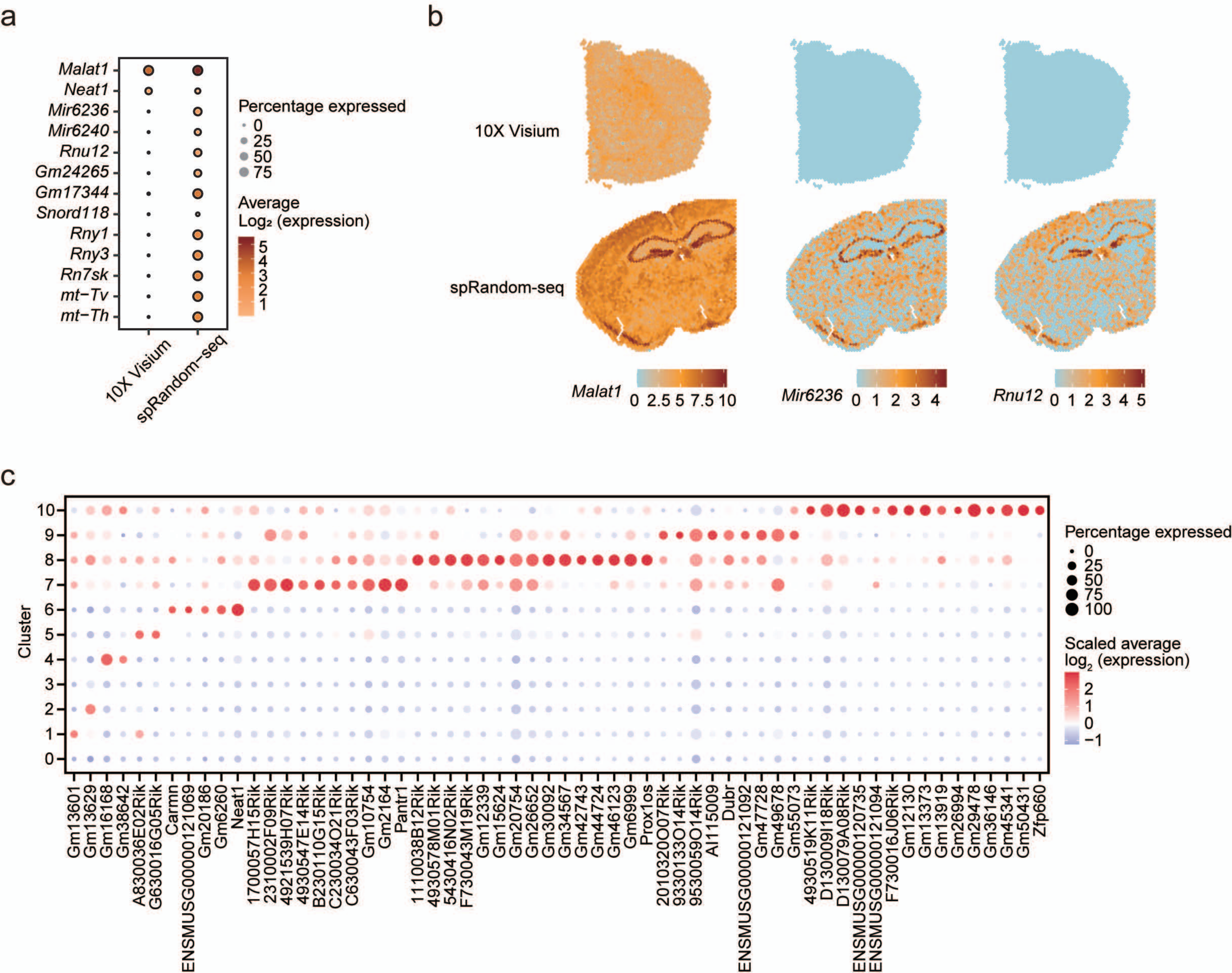
a
Malat1
Neat1
Mir6236
Mir6240
Rnu12
Gm24265
Gm17344
Snord118
Rny1
Rny3
Rn7sk
mt−Tv
mt−Th
10X Visium
spRandom−seq
Percentage expressed
0
25
50
75
Average
Log2 (expression)
5
4
3
2
1
b
10X Visium
spRandom-seq
Malat1
0 2.5 5 7.5 10
Mir6236
0 1 2 3 4
Rnu12
0 1 2 3 4 5
c
Cluster
10
9
8
7
6
5
4
3
2
1
0
Gm13601
Gm13629
Gm16168
Gm38642
A830036E02Rik
G630016G05Rik
Carmn
ENSMUSG00000121069
Gm20186
Gm6260
Neat1
1700057H15Rik
2310002F09Rik
4921539H07Rik
4930547E14Rik
B230110G15Rik
C230034O21Rik
C630043F03Rik
Gm10754
Gm2164
Pantr1
1110038B12Rik
4930578M01Rik
5430416N02Rik
F730043M19Rik
Gm12339
Gm15624
Gm20754
Gm26652
Gm30092
Gm34567
Gm42743
Gm44724
Gm46123
Gm6999
Prox1os
2010320O07Rik
9330133O14Rik
9530059O14Rik
AI115009
Dubr
ENSMUSG00000121092
Gm47728
Gm49678
Gm55073
4930519K11Rik
D130009I18Rik
D130079A08Rik
ENSMUSG00000120735
ENSMUSG00000121094
F730016J06Rik
Gm12130
Gm13373
Gm13919
Gm26994
Gm29478
Gm36146
Gm45341
Gm50431
Zfp660
Percentage expressed
0
25
50
75
100
Scaled average
log2 (expression)
2
1
0
−1

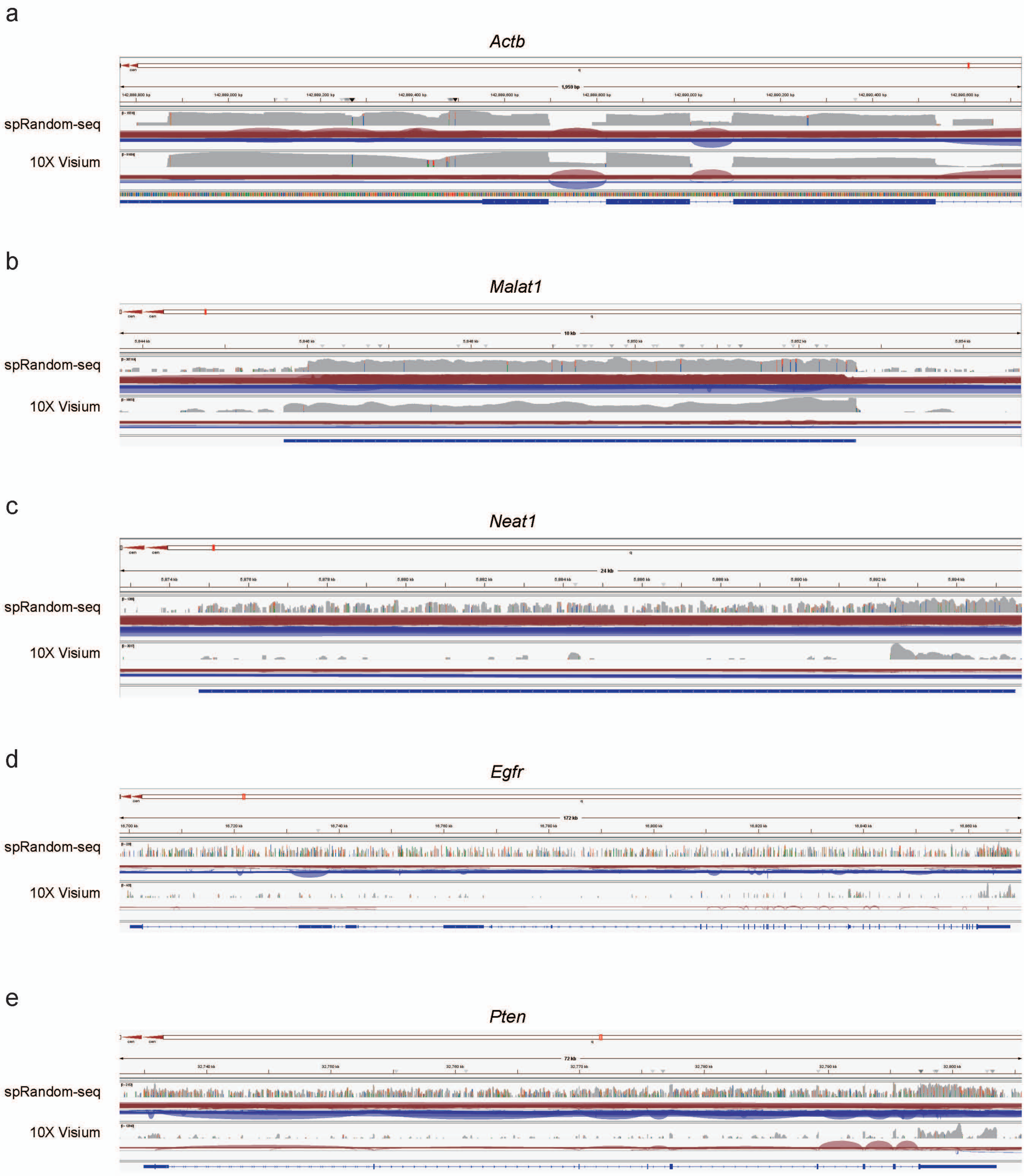
a
Actb
spRandom-seq
10X Visium
b
Malat1
spRandom-seq
10X Visium
c
Neat1
spRandom-seq
10X Visium
d
Egfr
spRandom-seq
10X Visium
e
Pten
spRandom-seq
10X Visium

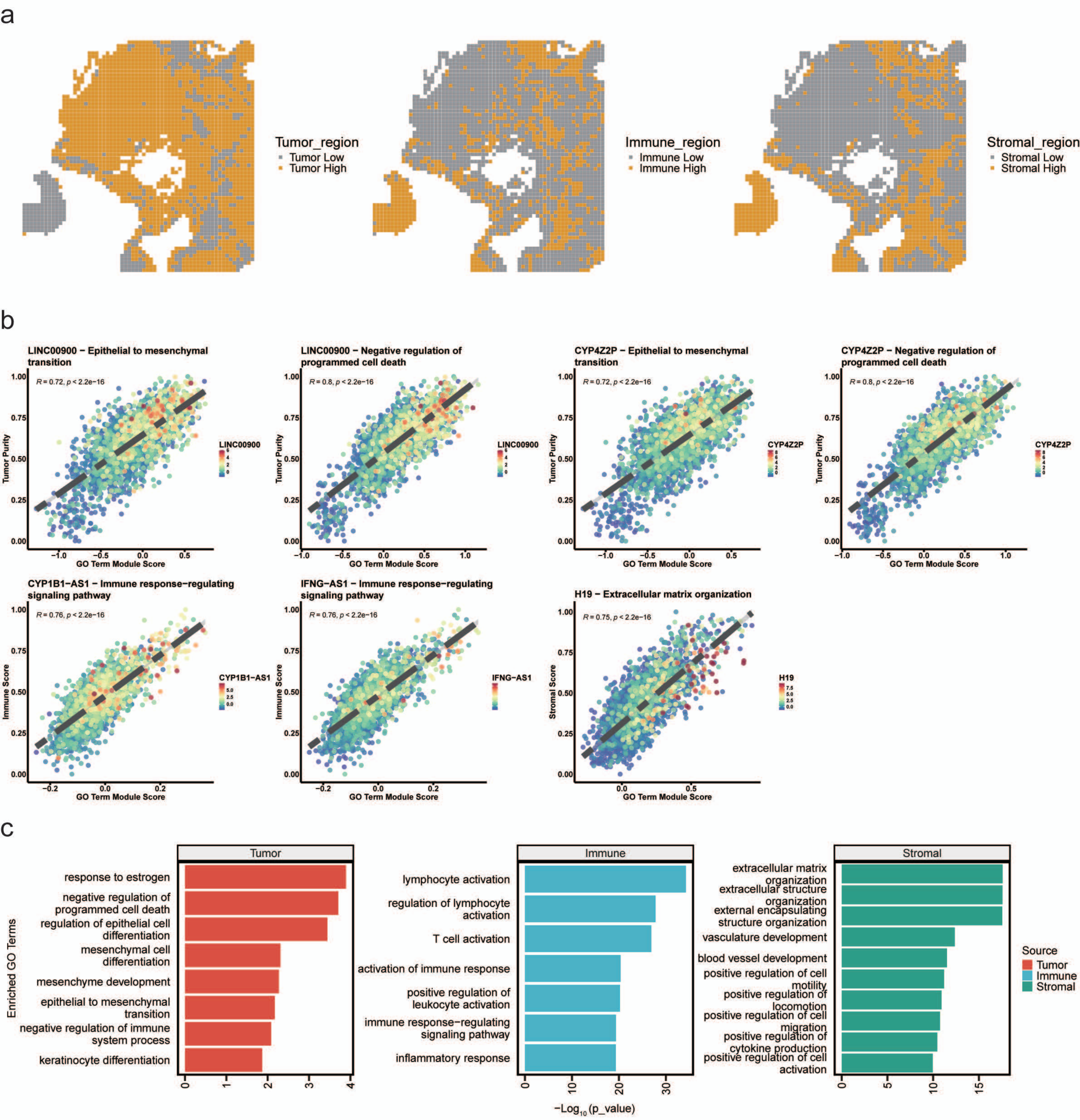
a
Tumor_region
Tumor Low
Tumor High
Immune_region
Immune Low
Immune High
Stromal_region
Stromal Low
Stromal High
b
LINC00900 − Epithelial to mesenchymal transition
R = 0.72, p < 2.2e−16
LINC00900 − Negative regulation of programmed cell death
R = 0.8, p < 2.2e−16
CYP4Z2P − Epithelial to mesenchymal transition
R = 0.72, p < 2.2e−16
CYP4Z2P − Negative regulation of programmed cell death
R = 0.8, p < 2.2e−16
Tumor Purity
GO Term Module Score
LINC00900
CYP4Z2P
CYP1B1−AS1 − Immune response−regulating signaling pathway
R = 0.76, p < 2.2e−16
IFNG−AS1 − Immune response−regulating signaling pathway
R = 0.76, p < 2.2e−16
H19 − Extracellular matrix organization
R = 0.75, p < 2.2e−16
Immune Score
Stromal Score
CYP1B1−AS1
IFNG−AS1
H19
c
Tumor
response to estrogen
negative regulation of programmed cell death
regulation of epithelial cell differentiation
mesenchymal cell differentiation
mesenchyme development
epithelial to mesenchymal transition
negative regulation of immune system process
keratinocyte differentiation
Immune
lymphocyte activation
regulation of lymphocyte activation
T cell activation
activation of immune response
positive regulation of leukocyte activation
immune response−regulating signaling pathway
inflammatory response
Stromal
extracellular matrix organization
extracellular structure organization
external encapsulating structure organization
vasculature development
blood vessel development
positive regulation of cell motility
positive regulation of locomotion
positive regulation of cell migration
positive regulation of cytokine production
positive regulation of cell activation
Enriched GO Terms
−Log10 (p_value)
Source
Tumor
Immune
Stromal

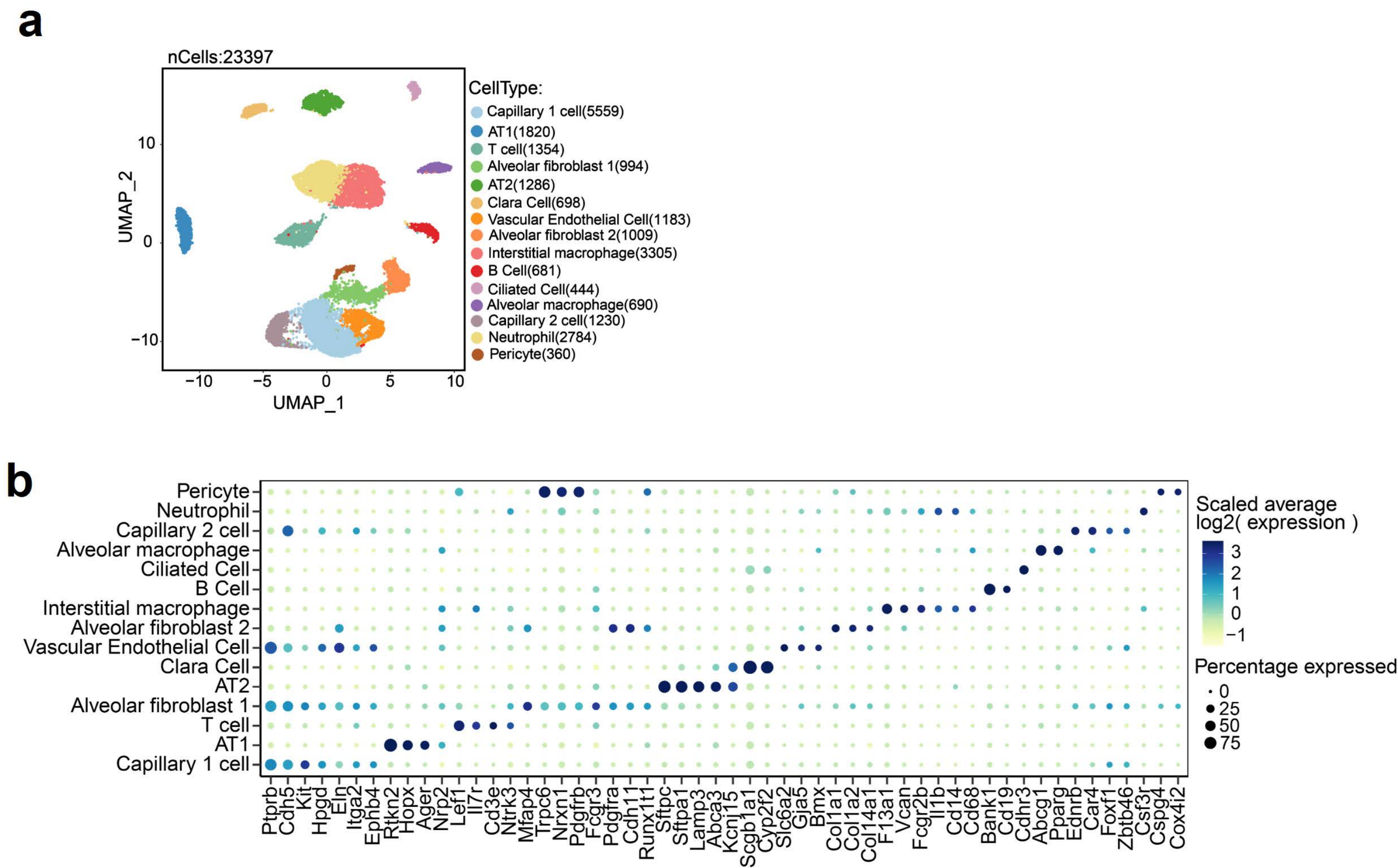
a
nCells:23397
UMAP_2
UMAP_1
10
0
−10
−10
−5
0
5
10
CellType:
Capillary 1 cell(5559)
AT1(1820)
T cell(1354)
Alveolar fibroblast 1(994)
AT2(1286)
Clara Cell(698)
Vascular Endothelial Cell(1183)
Alveolar fibroblast 2(1009)
Interstitial macrophage(3305)
B Cell(681)
Ciliated Cell(444)
Alveolar macrophage(690)
Capillary 2 cell(1230)
Neutrophil(2784)
Pericyte(360)
b
Pericyte
Neutrophil
Capillary 2 cell
Alveolar macrophage
Ciliated Cell
B Cell
Interstitial macrophage
Alveolar fibroblast 2
Vascular Endothelial Cell
Clara Cell
AT2
Alveolar fibroblast 1
T cell
AT1
Capillary 1 cell
Ptprb
Cdh5
Kit
Hpgd
Eln
Itga2
Ephb4
Rtkn2
Hopx
Ager
Nrp2
Lef1
Il7r
Cd3e
Ntrk3
Mfap4
Trpc6
Nrxn1
Pdgfrb
Fcgr3
Pdgfra
Cdh11
Runx1t1
Sftpc
Sftpa1
Lamp3
Abca3
Kcnj15
Scgb1a1
Cyp2f2
Slc6a2
Gja5
Bmx
Col1a1
Col1a2
Col14a1
F13a1
Vcan
Fcgr2b
Il1b
Cd14
Cd68
Bank1
Cd19
Cdhr3
Abcg1
Pparg
Ednrb
Car4
Foxf1
Zbtb46
Csf3r
Cspg4
Cox4i2
Scaled average
log2( expression )
3
2
1
0
−1
Percentage expressed
0
25
50
75

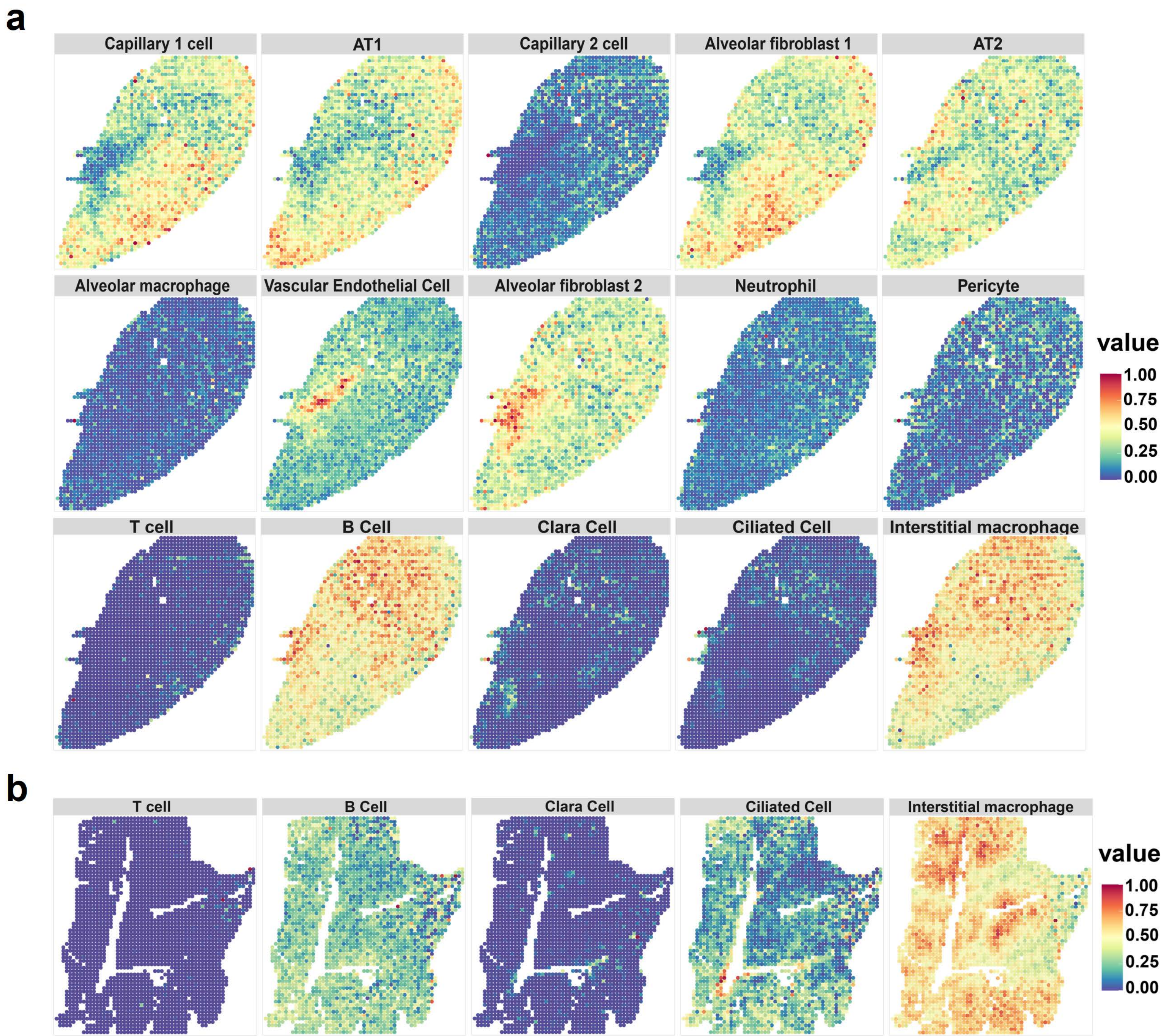

a
Capillary 1 cell
AT1
Capillary 2 cell
Alveolar fibroblast 1
AT2
Alveolar macrophage
Vascular Endothelial Cell
Alveolar fibroblast 2
Neutrophil
Pericyte
T cell
B Cell
Clara Cell
Ciliated Cell
Interstitial macrophage
value
1.00
0.75
0.50
0.25
0.00
b
T cell
B Cell
Clara Cell
Ciliated Cell
Interstitial macrophage
value
1.00
0.75
0.50
0.25
0.00

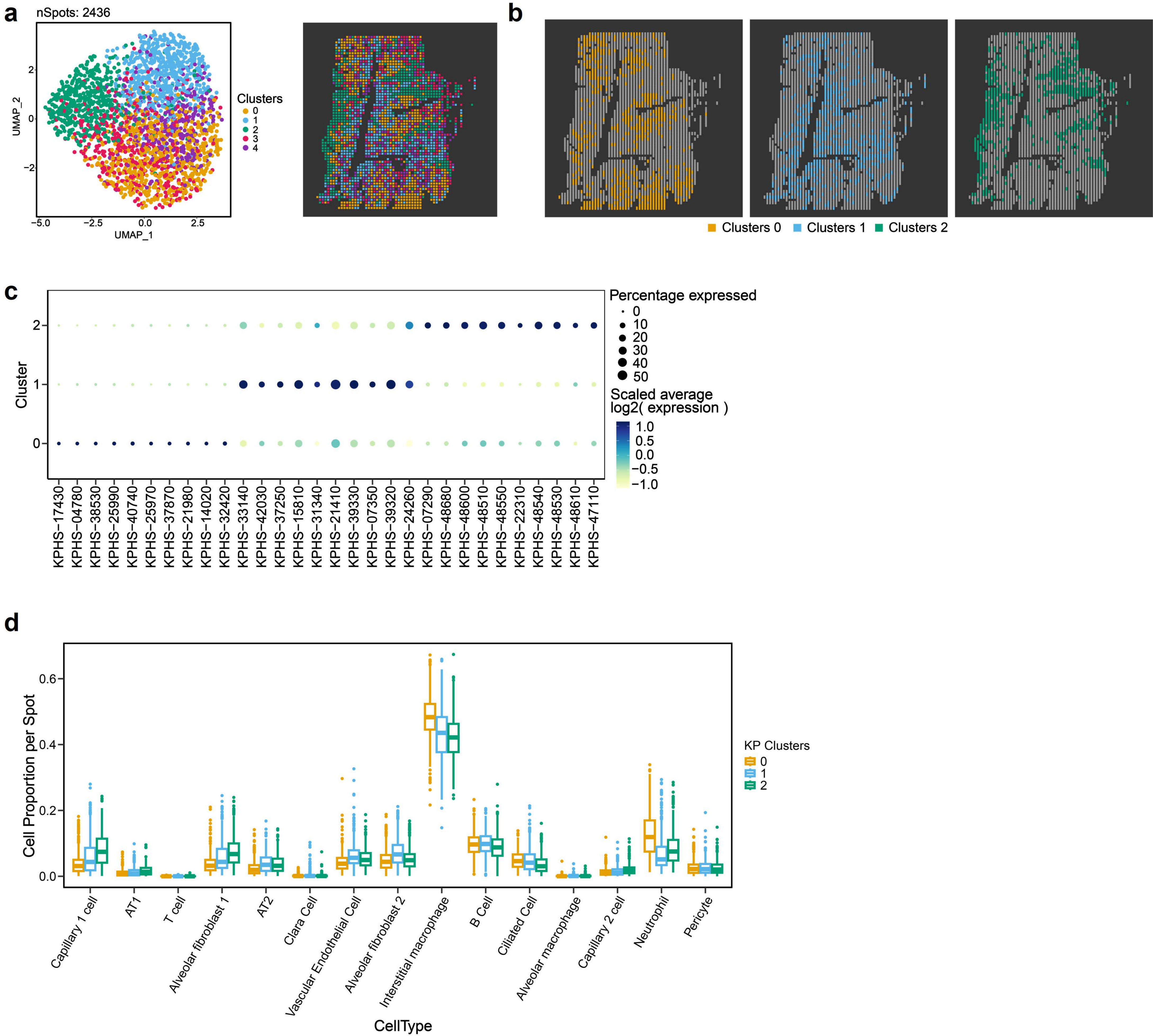
a
nSpots: 2436
UMAP_2
UMAP_1
−5.0
−2.5
0.0
2.5
2
0
−2
Clusters
0
1
2
3
4
b
Clusters 0
Clusters 1
Clusters 2
c
Cluster
2
1
0
Percentage expressed
0
10
20
30
40
50
Scaled average
log2( expression )
1.0
0.5
0.0
−0.5
−1.0
KPHS−17430
KPHS−04780
KPHS−38530
KPHS−25990
KPHS−40740
KPHS−25970
KPHS−37870
KPHS−21980
KPHS−14020
KPHS−32420
KPHS−33140
KPHS−42030
KPHS−37250
KPHS−15810
KPHS−31340
KPHS−21410
KPHS−39330
KPHS−07350
KPHS−39320
KPHS−24260
KPHS−07290
KPHS−48680
KPHS−48600
KPHS−48510
KPHS−48550
KPHS−22310
KPHS−48540
KPHS−48530
KPHS−48610
KPHS−47110
d
Cell Proportion per Spot
0.6
0.4
0.2
0.0
KP Clusters
0
1
2
Capillary 1 cell
AT1
T cell
Alveolar fibroblast 1
AT2
Clara Cell
Vascular Endothelial Cell
Alveolar fibroblast 2
Interstitial macrophage
B Cell
Ciliated Cell
Alveolar macrophage
Capillary 2 cell
Neutrophil
Pericyte
CellType

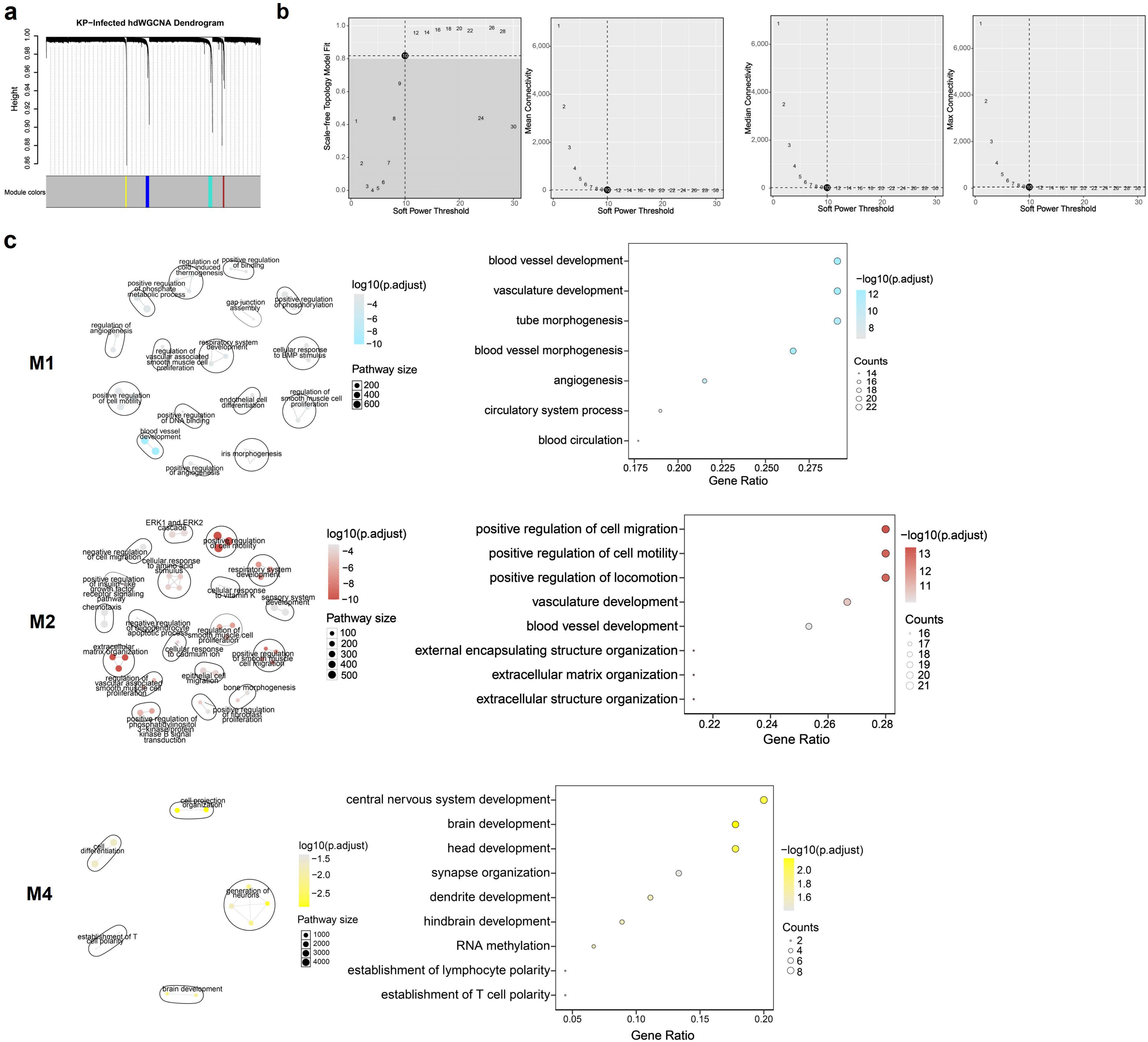

a
KP-Infected hdWGCNA Dendrogram
Height
Module colors
b
Scale-free Topology Model Fit
Mean Connectivity
Median Connectivity
Max Connectivity
Soft Power Threshold
c
M1
log10(p.adjust)
Pathway size
blood vessel development
vasculature development
tube morphogenesis
blood vessel morphogenesis
angiogenesis
circulatory system process
blood circulation
Gene Ratio
−log10(p.adjust)
Counts
M2
positive regulation of cell migration
positive regulation of cell motility
positive regulation of locomotion
vasculature development
blood vessel development
external encapsulating structure organization
extracellular matrix organization
extracellular structure organization
M4
central nervous system development
brain development
head development
synapse organization
dendrite development
hindbrain development
RNA methylation
establishment of lymphocyte polarity
establishment of T cell polarity

**Supplementary Fig. 1 | The comparison of different types of random primers**. **a**, The illustration of different primers. **b**, The CT values between various primers. There was no significant statistical difference among these primers.

**Supplementary Fig. 2 | The overview of CRISPR-based rRNA depletion**. **a**, The Cas9-based rRNA depletion and sequencing. **b**, The bases distributions of pre and after rRNA depletion and the pearson's correlation test on the expression of all genes, excluding rRNAs, pre and after rRNA depletion. **c**, The replication of CRISPR-based rRNA depletion for another section.

**Supplementary Fig. 3 | The RIN scores for mouse FFPE brain and heart.**

**Supplementary Fig. 4 | Spatial maps of the rest cell-type for spRandom-seq in FFPE mouse brain section.**

**Supplementary Fig. 5 | Validation and benchmark of spRandom-seq using a FFPE mouse heart section**. **a**, Spatial maps of classic cell-type for spRandom-seq in FFPE mouse heart section. **b**, Read coverage along the gene body by spRandom-seq in FFPE mouse heart section. **c**, Spatial maps of classic cell-type markers genes for spRandom-seq in the FFPE mouse heart section. Spots in which the transcript was not detected are shown as yellow. Green scale indicates log-normalized expression.

**Supplementary Fig. 6 ｜Detection of ncRNAs for spRandom-seq dataset**. **a**, Detection of coding and ncRNAs between 10X Visium and spRandom-seq workflows. Color scale shows average log-normalized UMI counts. Dot size shows the percentage of spots in which each RNA was detected. **b**, Spatial maps of coding and noncoding transcripts for 10X Visium and spRandom-seq datasets. Spots in which the transcript was not detected are shown as yellow. Color scale indicates log-normalized expression. **c**, Dotplot of top differentially expressed ncRNAs in each cluster based on the unsupervised clustering with only ncRNAs.

**Supplementary Fig. 7 ｜The IGV track for five genes from 10X Visium (fresh frozen mouse coronal brain) and spRandom-seq (FFPE mouse coronal brain): Actb, Malat1, Neat1, Egfr and Pten.**

**Supplementary Fig. 8 ｜Spatial ncRNA regulation in the clinical FFPE breast cancer section. a**, Spatial pattern of high or low tumor/immune/stromal region. Spots with high score (normalized value > 0.5) are shown as yellow. **b**, Top-ranked Enriched GO terms from the GO enrichment analysis performed on the differential expression protein-coding genes that showed positive correlations with the top upregulated ncRNAs in high tumor/immune/stromal regions respectively. **c**, Pearson's correlation tests between significant differential expression ncRNAs and signaling pathways.

**Supplementary Fig. 9 ｜The single-nuclear RNA sequencing data of the normal and infected mouse lungs using snRandom-seq. a**, UMAP plot of the integrated results of the normal and infected samples. Colored by 15 cell types identified. **b**, Dot plot of the average expressions of gene markers in each of the 15 cell types.

**Supplementary Fig. 10 ｜Spatial distribution maps for spRandom-seq in normal (a) and infected (b) samples.**

**Supplementary Fig. 11 ｜Spatial and functional heterogeneity of *K. pneumoniae* subpopulations in infected samples revealed by spRandom-seq. a**, UMAP projection and the spatial distribution of K. pneumoniae subpopulations. **b**, Spatial mappings of cluster 0, 1 and 2. **c**, Mean expression levels of top 10 DEGs for cluster 0, 1 and 2 of *K. pneumoniae*. **d**, Box plots of the distribution of cell type proportions at each spot within K. pneumoniae clusters 0, 1, and 2.

**Supplementary Fig. 12 | Analysis of gene co-expression modules in infected sample. a**, Dendrogram of gene co-expression modules, color-coded as Turquoise (M1), Brown (M2), Blue (M3), and Yellow (M4), identified through high-dimensional weighted gene co-expression network analysis. **b**, Optimal soft threshold selection in hdWGCNA. **c**, Gene Ontology (GO) pathway Enrichment Analysis of M1, M2 and M4 hub genes.

**Table S1. Comparison between spRandom-seq and other methods**

| **Comparison between 10X Visium and spRandom-seq (Human BRCA sample)** | | | | |
|---|---|---|---|---|
| Source | Sample Type | Sequencing saturation (%) | Mean Uniquely Mapped reads per spot | Valid barcode (%) |
| 10X Visium BRCA-Tumor (FF) | Fresh/Frozen | 72.1 | 41509 | 97.4 |
| spRandom-seq BRCA-Tumor (FFPE) | FFPE | 32.9 | 47153 | 99 |
| Source | Spots under tissue | Total genes detected | Median UMIs per spot | Median genes per spot |
| 10X Visium BRCA-Tumor (FF) | 4898 | 24387 | 9720 | 3654 |
| spRandom-seq BRCA-Tumor (FFPE) | 2682 | 38170 | 19759 | 4309 |
